# Cross-modal transfer enables zero-shot prediction of drug responses across cell types

**DOI:** 10.64898/2026.10.27.762107

**Authors:** Cheng Luo, Miguel Hu, Yahong Wu, Danni Guo, Songmao Wang, Qing Rex Lyu, Xiao Liu

**Affiliations:** Institute of Biopharmaceutical and Health Engineering, Tsinghua Shenzhen International Graduate School, Tsinghua University, Shenzhen 518055, China; Department of Cardiology, Chongqing Academy of Medical Science, Chongqing General Hospital, Chongqing University, Chongqing 401147, China

## Abstract

Genetic perturbation responses capture the functional consequences of gene-level interventions, providing functional information for drug-response prediction beyond baseline gene expression. Within the same cellular context, genetic and chemical perturbations engage shared regulatory networks, providing a biological basis for transferring response information between perturbation modalities. Here we introduce **GRAFT**, a framework that combines genetic-to-chemical transfer with cross-context prediction to infer drug-induced transcriptional responses without drug measurements in the target context. Across benchmarked LINCS and external contexts, GRAFT improved response correlation, directional agreement and magnitude reconstruction over competing methods, particularly for drugs with context-dependent responses. **GRAFT-Adapt** further refines predictions of unmeasured responses using a small number of drug measurements from the target context. **GRAFT-Atlas** spans 13 cellular contexts and 7,856 drug catalog entries, comprising 102,128 predictions for drug–context combinations. The atlas captured recognizable pharmacological neighborhoods, shared functional programs, context-dependent transcriptional changes and structural–functional associations supported by molecular docking. Across ten iPSC lines from distinct donors, GRAFT identified candidate inter-donor differences, with functional support for selected candidates from measured genetic perturbation responses. Together, GRAFT enables perturbation-informed prediction and comparison of drug responses across cellular contexts.

## Introduction

The same drug can induce different transcriptional changes across cellular contexts^1,2^. Predicting responses in a new context therefore requires both transferable pharmacological information and a representation of how the target cells respond. Large-scale perturbation atlases have linked compounds and genetic interventions to downstream transcriptional changes, but the space of drug–context combinations remains difficult to measure comprehensively^3,4^. Genetic perturbation measurements provide a direct readout of cellular responses to intervention, providing functional information about the target context^5^. Genetic and chemical interventions can affect interconnected regulatory pathways and converge on shared transcriptional programs, suggesting that genetic responses may provide target-context functional information for drug-response models trained in other contexts^3,6^.

Achieving this transfer requires combining genetic-to-chemical prediction with adaptation across cellular contexts. Methods such as XPert and State integrate unperturbed expression with perturbation information to predict cellular responses^7,8^. However, unperturbed expression describes the baseline state, whereas the response to an intervention also depends on context-specific regulation. Similar baseline expression therefore does not necessarily imply similar perturbation responses^9^. UniPert-G2CP demonstrates the value of genetic-to-chemical transfer through unified perturbation representations, with transfer models trained within individual cell lines^6^. HyperMap uses a small number of target perturbation responses to adapt across contexts, but adapts genetic and chemical perturbations separately^9^. Cross-context benchmarks further emphasize the importance of cellular context as an important direction for improving generalization^10^. These studies leave open how genetic responses measured in target context can guide drug-response prediction without target drug measurements. Drug responses across source contexts provide references for transferable pharmacological effects, while differences between target and source genetic responses offer information for context specific corrections.

Here we present GRAFT, a framework for genetic-to-chemical transfer that enables zero-shot prediction of transcriptional responses without drug measurements in the target cellular context. GRAFT learns relationships between genetic and drug responses across multiple source cell contexts and aggregates differences between target and source genetic responses into a conditioning state. This state modulates a shared prediction readout to adjust source-averaged responses for individual drugs, allowing the same genetic dataset to support predictions across multiple drugs.

When limited target drug measurements are available, GRAFT-Adapt uses their prediction residuals to refine predictions for unmeasured drugs. Across five LINCS cancer cell lines and five external contexts from public datasets, GRAFT outperformed multiple methods in response correlation and magnitude reconstruction. Genetic perturbation budget analyses further evaluated prediction and adaptation performance when limited target genetic measurements were available. We also constructed GRAFT-Atlas across 13 cellular contexts, expanding drug-response coverage to examine pharmacological relationships, shared transcriptional programs and context-specific responses.

## Results

### GRAFT links genetic responses to drug-response prediction

We developed GRAFT (Genetic Response-guided Adaptive Forecasting of drug-induced Transcriptomes) to predict drug responses across cellular contexts using genetic responses from the target cell line. During training, source cell lines alternate as prediction targets, learning to use genetic-response differences to correct drug-response predictions (**Fig. 1a**). UniPert supplies genetic perturbation identity representations incorporating protein sequence information^6^. For each genetic perturbation, the model combines this representation with its mean response across source contexts. A parallel branch encodes the difference between the target genetic response and this source reference. Cross-moment pooling summarizes their paired variation into a conditioning state shared across drug queries. Each query combines drug identity with the mean response to that drug across source contexts, termed the source mean. A hypernetwork converts the conditioning state into feature scaling and shifting parameters applied through feature-wise linear modulation (FiLM)^11^.

**Figure 1.**
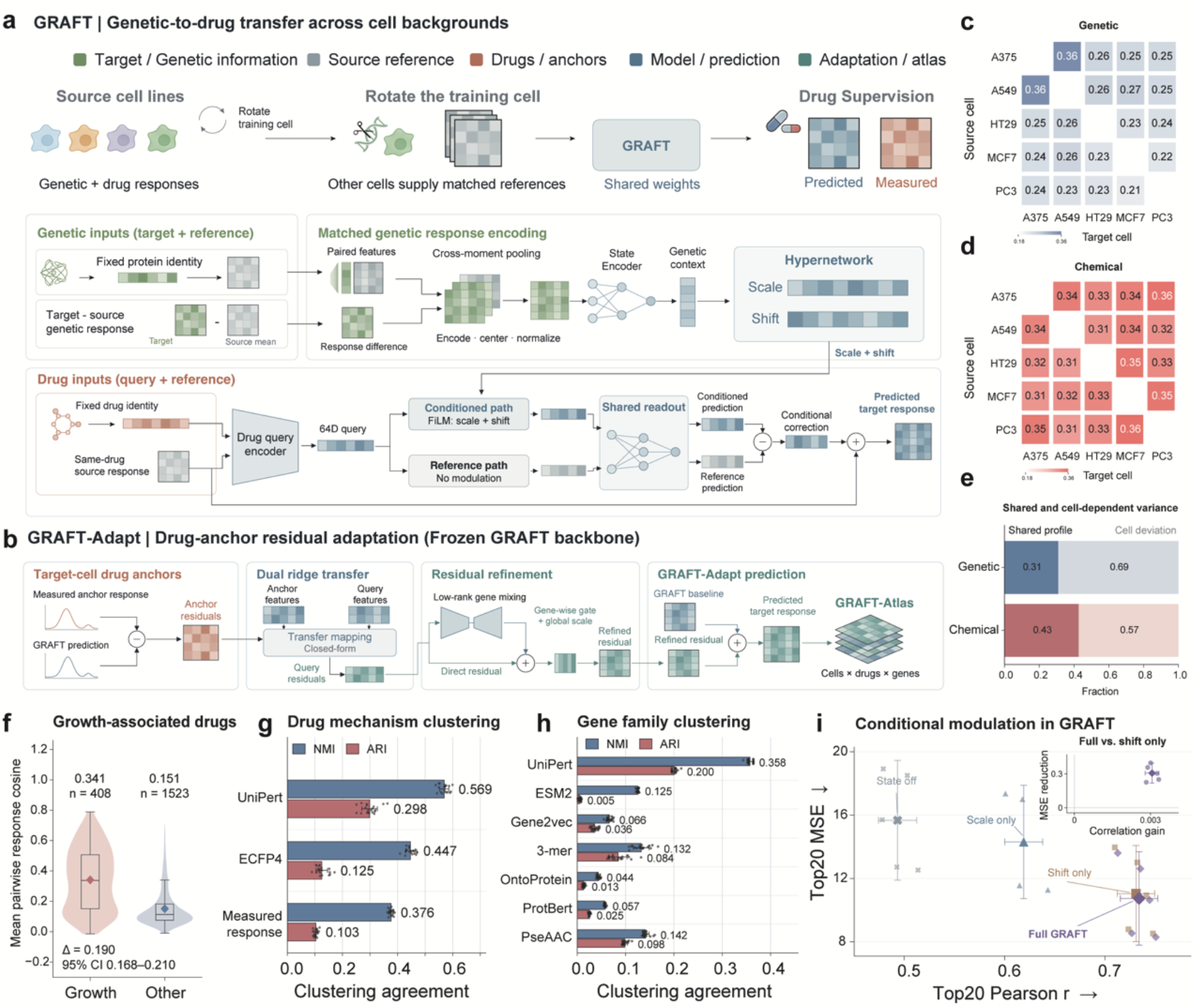
GRAFT architecture and perturbation-response organization. **a**,**b**, Genetic-response conditioning for drug prediction (a) and drug-anchor adaptation with a frozen GRAFT backbone (b). **c**,**d**, Directed source-to-target Pearson correlations for 4,156 genetic perturbations (c) and 3,220 drugs (d), calculated over each source cell’s 20 genes with the largest absolute responses. **e**, Fractions of the total response sum of squares attributed to shared profiles and cell-specific deviations across 976 genes. **f**, Cross-cell response cosine similarity for 408 growth-associated and 1,523 other annotated drugs. Boxes show medians and interquartile ranges (IQRs); whiskers extend to observations within 1.5×IQR; diamonds indicate means. The mean-difference interval is a 95% drug-bootstrap confidence interval (CI). **g**,**h**, Clustering agreement with drug mechanisms (527 drugs, 27 classes; g) and gene families (524 genes, six classes; h), measured by normalized mutual information (NMI) and adjusted Rand index (ARI). Bars show mean ± s.d. across 20 clustering seeds. **i**, Modulation ablations with trained model parameters held fixed. Small symbols show five cell-line means over three model seeds; large symbols and whiskers show means and t-based 95% CIs. The inset compares full GRAFT with shifting alone. MSE, mean squared error.

Conditioned and reference features pass through the same readout and the difference between their outputs is used to correct the source mean. Reusing the conditioning state across drug queries allows one set of target genetic measurements to support predictions for multiple drugs.

When a small number of drug-response measurements are available in the target cell line, GRAFT-Adapt uses them to calibrate the base prediction. With the GRAFT backbone frozen, GRAFT-Adapt computes residuals between measured anchor-drug responses and base predictions, then uses dual-form ridge regression on conditioned drug features to estimate corrections for unmeasured drugs. Low-rank gene mixing and gating refine the estimated correction which is added to the base prediction alongside a direct residual path (**Fig. 1b**). We also integrated available genetic and drug perturbation data across 13 cellular contexts to construct GRAFT-Atlas. Using a common drug catalog, GRAFT-Atlas organizes predicted transcriptional responses across contexts for comparison of pharmacological relationships, shared functional programs and context-dependent differences.

Measured responses supported combining shared references with context-dependent correction. Across A375, A549, HT29, MCF7 and PC3, drug responses were more correlated across contexts than genetic responses (mean Top20 Pearson *r*, 0.332 versus 0.255; **Fig. 1c,d**). The same contrast appeared with broader gene sets and principal-component summaries (**Extended Data Fig. 1a,b**). Cell-specific deviations accounted for more than half of the variance in both modalities (**Fig. 1e**). Source drug responses may therefore provide a starting point for prediction, while genetic responses from the target cell line could inform context-specific corrections. Growth-associated drugs showed more consistent responses than several receptor ligands (**Fig. 1f** and **Extended Data Fig. 1c**,**e**). Drugs with more shared responses included inhibitors of CDK, mTOR, HDAC, topoisomerases and EGFR, such as alvocidib, palbociclib, vorinostat and NVP-BEZ235, whereas budesonide and fulvestrant were more context dependent. The context dependence of these steroid-receptor modulators is consistent with the role of chromatin in steroid-receptor regulation^12^. These observations support prediction that retains shared pharmacological responses while allowing context-dependent differences.

Perturbation identity representations also captured pharmacological relationships. UniPert organized drugs and genes by mechanism of action and pharmacological family (**Fig. 1g,h**). Within-class similarity showed larger effect sizes for UniPert drug representations than for molecular fingerprints. PI3K inhibitors remained close in this representation despite dispersed chemical fingerprints (**Extended Data Fig. 1d**,**f**).

Genetic–drug response correlations were higher for target-linked inhibitors and antagonists than for agonist-like drugs across the five cell lines (**Extended Data Fig. 1g**,**h**). These correlations also exceeded those of structure-matched non-target pairs, indicating that chemical similarity alone did not explain the correspondence (**Extended Data Fig. 1i**).

Ablations with the trained model held fixed assessed the contribution of the conditioning state. Disabling it reduced Top20 Pearson *r* from 0.734 to the value for the source mean of 0.493. Feature shifting retained most predictive capacity, while scaling further improved correlation and reduced magnitude error (**Fig. 1i**). Within the 32-dimensional state, retaining the 16–24 largest-magnitude components preserved most performance and was more effective than random retention (**Extended Data Fig. 2a**,**b**). Greater capacity, attention, gene graphs and query interactions did not consistently improve predictions (**Extended Data Fig. 2c**,**e**). Similar performance across set encoders supported the compact cross-moment interface.

**Figure 2.**
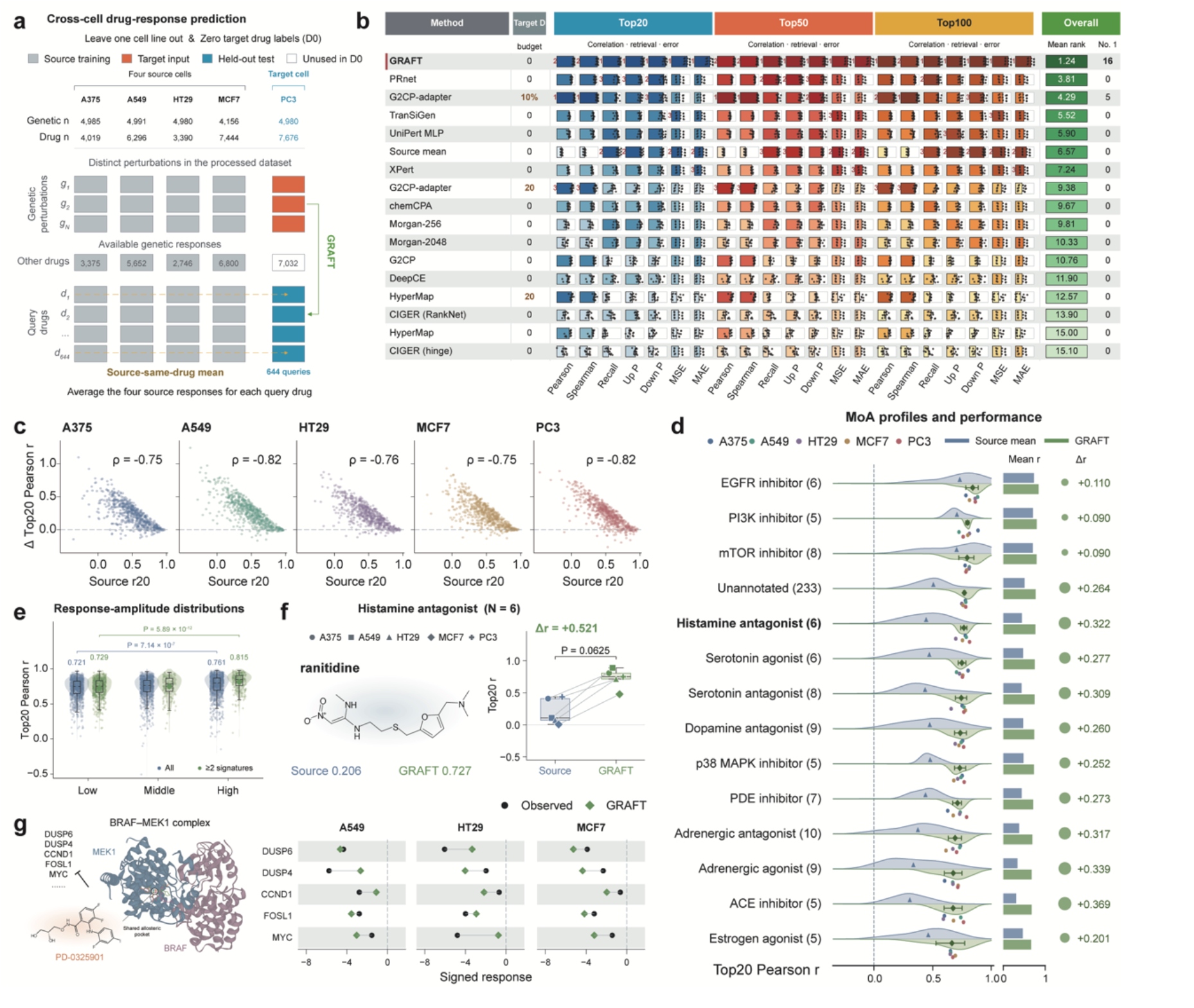
Cross-cell drug prediction and pharmacological readouts. **a**, Five-cell line leave-one-out D0 evaluation of 644 drugs across 976 genes, with no target-drug measurements used for prediction. **b**, GRAFT and 16 comparator configurations evaluated across 21 metrics. Bars show normalized performance; dots denote five cells × three seeds. Overall reports mean rank and first-place counts; Target D denotes target-drug budget. **c**, Source-mean correlation versus GRAFT gain for 641 drugs per cell; *ρ* denotes Spearman correlation. **d**, Performance by mechanism of action (MoA). Blue/green distributions and bars represent source/GRAFT scores; triangles/diamonds denote means, colored dots indicate cells and bubble area encodes gain. Counts denote drugs. **e**, Scores across within-cell measured-response-amplitude tertiles, using all records or those with ≥2 signatures (3,205/1,018 drug–cell pairs). *P* values are from cell-adjusted, drug-cluster-robust t tests, with Holm correction. **f**, Ranitidine, selected for the largest mean gain among six histamine antagonists. Lines connect five cell pairs; the displayed *P* value is from an unadjusted two-sided exact paired sign-flip test. **g**, PD-0325901 structural reference (PDB 7M0X) and measured/predicted responses of five genes in three cells. Intervals in d,e are 95% drug-bootstrap confidence intervals. Boxes in e,f indicate medians/interquartile ranges (IQRs), with whiskers extending to observations within 1.5×IQR. Panels d–g use three-seed prediction ensembles. Top20 *r* is Pearson correlation over the 20 genes with largest absolute measured responses; Δr = GRAFT − source mean.

Input permutation experiments further examined which relationships among genetic-response features were necessary for prediction. Permuting raw genetic responses while preserving source-reference correspondence had little effect. Disrupting correspondence among genetic residuals, perturbation identities and source-reference features reduced performance (**Extended Data Fig. 2d**). In residual ridge-regression probes fitted using measured target drugs excluded from the queries, GRAFT-derived drug features outperformed the original identity representations (**Extended Data Fig. 2f**,**g**).

### GRAFT improves drug-response prediction across LINCS cell lines

We evaluated cross-context transfer in five LINCS cancer cell lines, training on four and holding out the fifth as the target^3^. Each fold predicted 644 query drugs across 976 genes. In the D0 setting, GRAFT received target genetic responses and source responses to the same drugs, while target drug measurements were reserved for evaluation (**Fig. 2a**).

GRAFT-D0 achieved a mean Top20 Pearson *r* of 0.734, compared with 0.493 for the source mean and 0.689 for PRnet, the strongest D0 comparator. The correlation advantage held across all five targets and three seeds (**Extended Data Fig. 3a**). Against 16 comparator configurations evaluated under different target-drug budgets, GRAFT ranked first on 16 of 21 metrics (**Fig. 2b**). Comparators included PRnet, DeepCE, CIGER, TranSiGen and chemCPA^4,13-16^. Improvements extended to Top50 and Top100 windows, magnitude errors and strong-response gene retrieval (**Extended Data Fig. 3b**).

**Figure 3.**
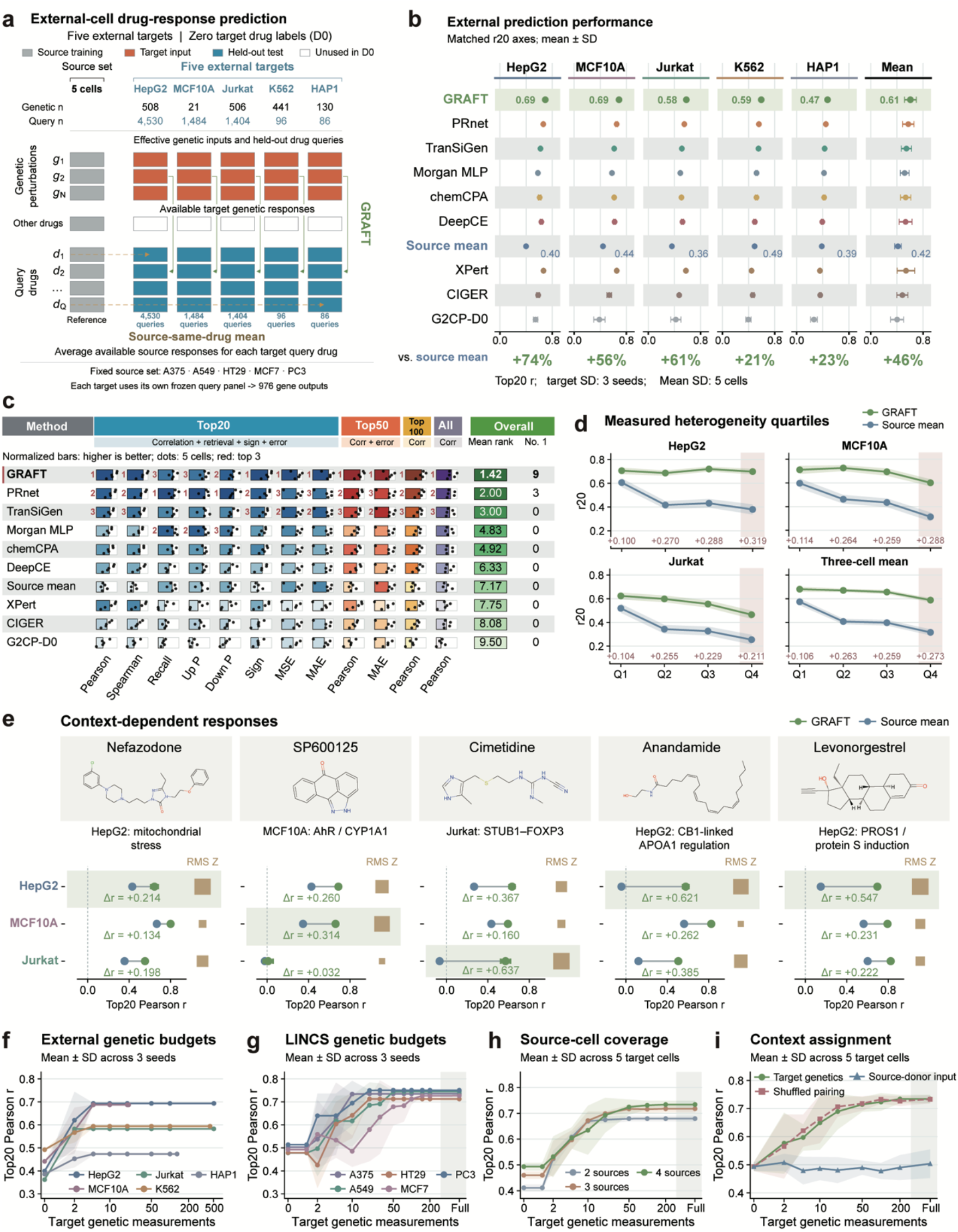
External prediction and genetic-input requirements. **a**, D0 evaluation in five external contexts using five LINCS sources and target genetic responses across 976 genes. Counts indicate genetic inputs and drug queries. **b**, Top20 Pearson correlations for ten configurations. Points for individual targets show means ± s.d. across 3 seeds; the summary column shows the mean ± s.d. across five target-context means. Percentages indicate improvement over the source mean. **c**, Twelve-metric comparison with normalized bars (higher is better), five target means and ranks from raw macro-averages. **d**, Performance for 650 drugs shared by HepG2, MCF10A and Jurkat, grouped by measured heterogeneity quartiles (1 − mean pairwise full-profile Pearson correlation). Shading indicates 95% drug-bootstrap confidence intervals. **e**, Five highest-quartile drugs: correlations show three-seed mean ± s.d.; gold square area encodes measured root-mean-square (RMS) Z-score across the three contexts within each drug. **f**,**g**, Genetic-input budgets in external (f) and LINCS (g) contexts; curves and shading show three-seed mean ± s.d. after averaging input subsamples. **h**, Comparison of two, three and four source cells. **i**, Target, shuffled-pairing and source-donor genetic inputs. In **h**,**i**, curves and shading show the mean ± s.d. across five target contexts, after averaging over source combinations, seeds and subsamples within each context.

GRAFT-D0 also exceeded the prediction correlations of G2CP and HyperMap given 20 non-query target drugs^6,9^. With approximately 10% of the target drug catalog, G2CP achieved slightly higher values on some correlation metrics, but GRAFT-D0 retained lower magnitude errors and better strong-response gene recall (**Extended Data Fig. 3c**,**d**). About 93% of drug–cell-line pairs improved over the source mean. This broad distribution showed that the improvements were not driven by a small number of high-gain examples. We defined correlation gain as GRAFT’s prediction correlation minus the correlation of the source mean. Drugs predicted poorly by the source mean generally showed larger gains. Correlation of the source mean and correlation gain were inversely associated across targets (Spearman *ρ*, −0.82 to −0.75; **Fig. 2c**).

Performance varied with pharmacological class and measured response strength. EGFR, PI3K and mTOR inhibitors retained high correlations, whereas several receptor-ligand classes, including histamine antagonists, showed larger gains over weaker source predictions (**Fig. 2d**). Mean Top20 Pearson *r* was higher in the high-than low-response-amplitude tertile (0.761 versus 0.721). This difference was larger among profiles supported by at least two signatures (0.815 versus 0.729; **Fig. 2e**). Among histamine antagonists, ranitidine, a competitive histamine H2-receptor antagonist, showed a particularly large improvement. For ranitidine, GRAFT improved mean Top20 Pearson *r* from 0.206 to 0.727, with gains across all five cell lines (**Fig. 2f**). Results for A-66, tropisetron and amisulpride provided additional examples of improved predictions across different pharmacological classes (**Extended Data Fig. 3e**).

PD-0325901 provided a gene-level example of recognizable pharmacology. In A549, HT29 and MCF7, GRAFT recovered the measured downregulation of *DUSP6, DUSP4, CCND1, FOSL1* and *MYC* (**Fig. 2g**). These markers are linked to MEK– ERK-dependent transcription and responses to pathway inhibition^17^. GRAFT thus recovered coordinated transcriptional repression downstream of MEK inhibition across these cellular contexts.

### GRAFT generalizes to external contexts with limited genetic measurements

We evaluated GRAFT in five external cellular contexts, HepG2, MCF10A, Jurkat, K562 and HAP1, using public genetic and drug perturbation data^3,5,18-20^. These targets provided 21–508 usable genetic inputs, compared with thousands per LINCS context. Their limited and uneven coverage therefore tested cross-context transfer from sparse target genetic measurements, a common setting when applying models to new cellular contexts (**Fig. 3a**). Drug responses matching the source catalog supplied 4,530 queries in HepG2, 1,484 in MCF10A, 1,404 in Jurkat, 96 in K562 and 86 in HAP1. The equally weighted mean Top20 *r* across contexts reached 0.606, compared with 0.416 for the source mean and 0.575 for the strongest macro-average comparator. GRAFT achieved the highest Top20 correlation among the compared D0 models in every external target (**Fig. 3b**). Advantages extended to response direction, magnitude reconstruction and broader strong-response gene sets (**Fig. 3c; Extended Data Fig. 4a–e**).

**Figure 4.**
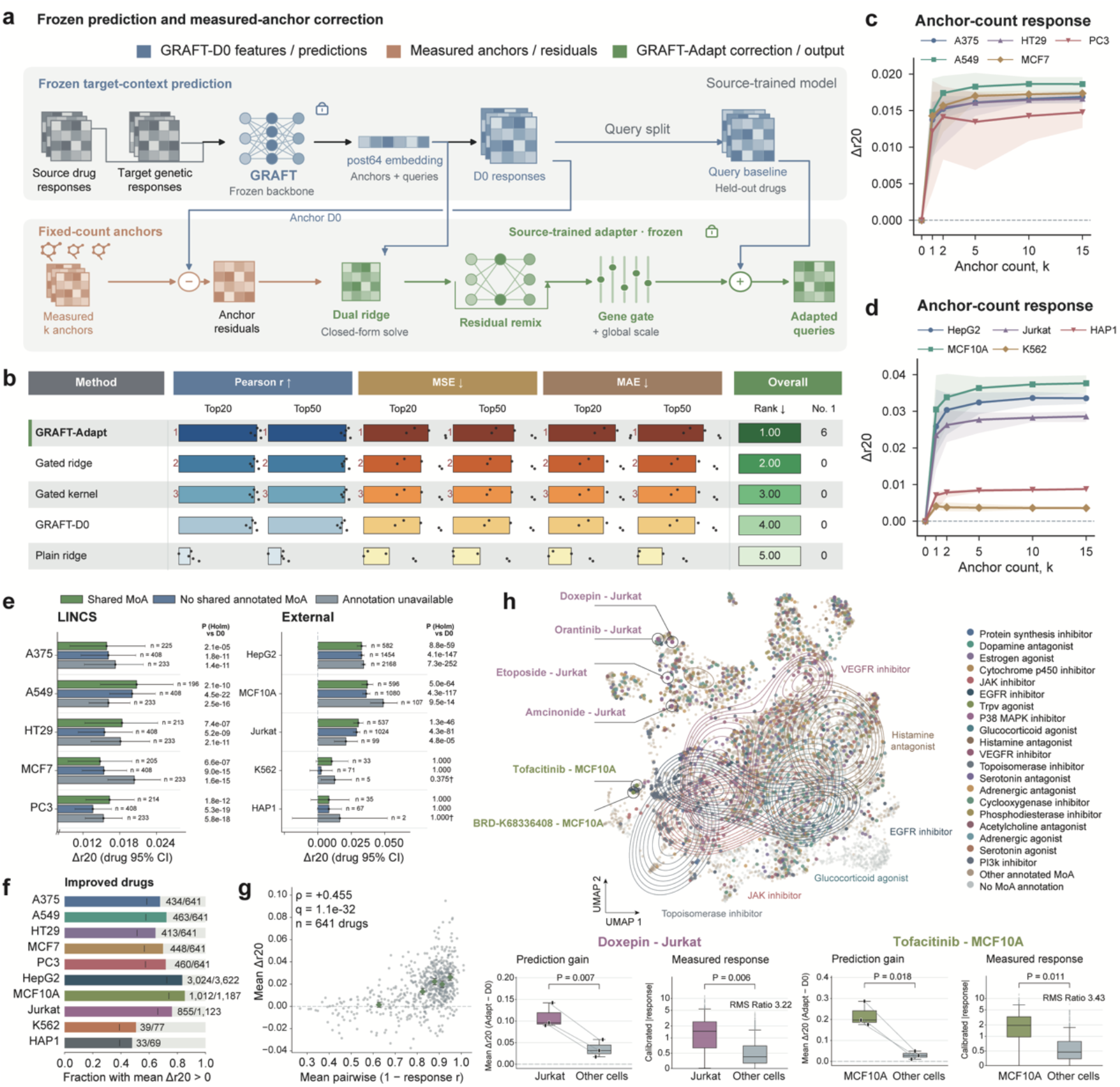
Drug-anchor adaptation across cellular contexts. **a**, Frozen-model correction using measured anchor-drug residuals. **b**, Five-method LINCS comparison across six post hoc-selected reconstruction metrics. Bars show normalized performance; dots denote five target means. **c**,**d**, Anchor-budget gains in internal (c) and external (d) contexts. Shading in c,d shows the empirical 95% range across 30 anchor lists after averaging three seeds. e, Ten-anchor gains stratified by whether query drugs share an annotated mechanism of action (MoA) with any anchor drug. Intervals show 95% drug-bootstrap confidence intervals; P values are from two-sided paired Wilcoxon tests, with Holm adjustment across 30 context–stratum comparisons. Counts denote unique drugs per stratum. **f**, Fractions of drugs with positive mean gains. **g**, Measured response heterogeneity versus mean gain for 641 LINCS drugs; diamonds show median heterogeneity and mean gain within quintiles, with 95% drug-bootstrap intervals. Spearman *ρ* and Benjamini–Hochberg-adjusted *q* are shown. **h**, Measured-response UMAP colored by MoA. Below, doxepin–Jurkat and tofacitinib–MCF10A are compared with the same drugs in other contexts. Gain comparisons use three paired seed means after averaging 30 lists (two-sided paired t tests). Response distributions contain calibrated measured responses across 976 genes per context; response *P* values are drug-reference empirical tail probabilities for focal/other-context RMS ratios (350 reference drugs per case). Displayed *P* values are unadjusted. Boxes indicate medians/interquartile ranges (IQRs); whiskers extend to observations within 1.5×IQR. Unless varied in **c**,**d**, ten anchors were used. Δr20 = GRAFT-Adapt − D0.

Drugs with greater measured cross-context differences generally showed larger improvements over the source mean. We grouped drugs measured in HepG2, MCF10A and Jurkat by quartiles of heterogeneity in their complete response profiles. Every quartile showed positive gains over the source mean. Mean correlation gains were 0.273 in the highest quartile and 0.106 in the lowest (**Fig. 3d**). Individual drug– context examples likewise showed larger gains where measured responses were stronger (**Fig. 3e**). Class-stratified analyses also showed large gains for glucocorticoid- and estrogen-receptor agonists in HepG2, MCF10A and Jurkat (**Extended Data Fig. 4f**). These results indicate that GRAFT’s advantage over the source mean was larger for drugs with more heterogeneous measured responses across contexts.

We next examined how prediction performance changed with the number of available target genetic perturbations. The genetic perturbation budget was the number of genetically perturbed targets whose responses in the target cell line were supplied as model inputs. Increasing the number of target genetic perturbations generally improved external prediction. With only 21 usable measurements, MCF10A reached a Top20 *r* of 0.687, exceeding the source mean of 0.441 (**Fig. 3f)**. Across the five LINCS cancer cell lines, approximately 200 inputs approached the mean correlation obtained with thousands of measurements (**Fig. 3g**). Increasing genetic inputs also improved broader-window correlations and gene retrieval while reducing magnitude errors in LINCS (**Extended Data Fig. 4g–j**). These results suggest that GRAFT can improve drug-response predictions without requiring extensive genetic perturbation measurements in every target context.

Transfer performance also depended on source-context coverage and whether the genetic inputs came from the target cells. Increasing the number of source cell lines improved prediction when all target genetic inputs were available (**Fig. 3h**).

Replacing target responses with source-cell responses failed to recover the same gains, whereas raw-response permutation preserving source-reference correspondence had little effect (**Fig. 3i**). Directly disrupting genetic residual correspondences reduced performance (**Extended Data Fig. 2d**). These controls indicated that transfer depended on a conditioning state formed from target genetic responses relative to source references.

### GRAFT-Adapt improves predictions with limited drug measurements

We next asked whether drug anchors could correct errors remaining after genetic conditioning. Anchor residuals were differences between measured target responses and GRAFT-D0 predictions. GRAFT-Adapt transferred these residuals to queries through dual-form ridge regression on conditioned drug features, followed by low-rank mixing, gene-wise gating and global scaling before addition to the base prediction. The GRAFT backbone and adapter parameters remained frozen; anchor measurements determined only the residual-mapping coefficients, while query drugs remained held out. We compared GRAFT-Adapt with GRAFT-D0 on the same queries across five internal cancer cell lines and five external contexts. We used repeated anchor sampling with matched fixed-count budgets, selecting 10 anchors for the primary comparison and using identical anchor sets across methods. This matched the number of target drug measurements used across contexts (**Fig. 4a**).

With identical anchor and query splits, GRAFT-Adapt ranked first against standard ridge, gated ridge and gated kernel regression on all six reported internal reconstruction metrics. These comprised correlation, mean squared error (MSE) and mean absolute error (MAE) within Top20 and Top50 windows (**Fig. 4b**). Most average improvement occurred with the first few anchors, after which gains plateaued.

Results were consistent across anchor sets and model seeds (**Fig. 4c,d**). Mean changes were also positive in K562 and HAP1. However, their small query panels and candidate anchor pools, derived from different datasets, may increase uncertainty in these gains.

Correction extended beyond pharmacological classes represented among the anchors. Across all five internal contexts and HepG2, MCF10A and Jurkat, queries sharing no annotated mechanism of action with anchors showed positive gains. Their drug-resampling intervals lay entirely above zero (**Fig. 4e**). These results suggest that the learned drug features support residual transfer beyond shared annotated mechanisms of action. Positive correlation gains over GRAFT-D0 occurred for approximately 64%–72% of drugs across the internal cancer cell lines. The corresponding proportions were approximately 76%–85% in the three larger external query panels, HepG2, MCF10A and Jurkat (**Fig. 4f; Extended Data Fig. 5a**).

**Figure 5.**
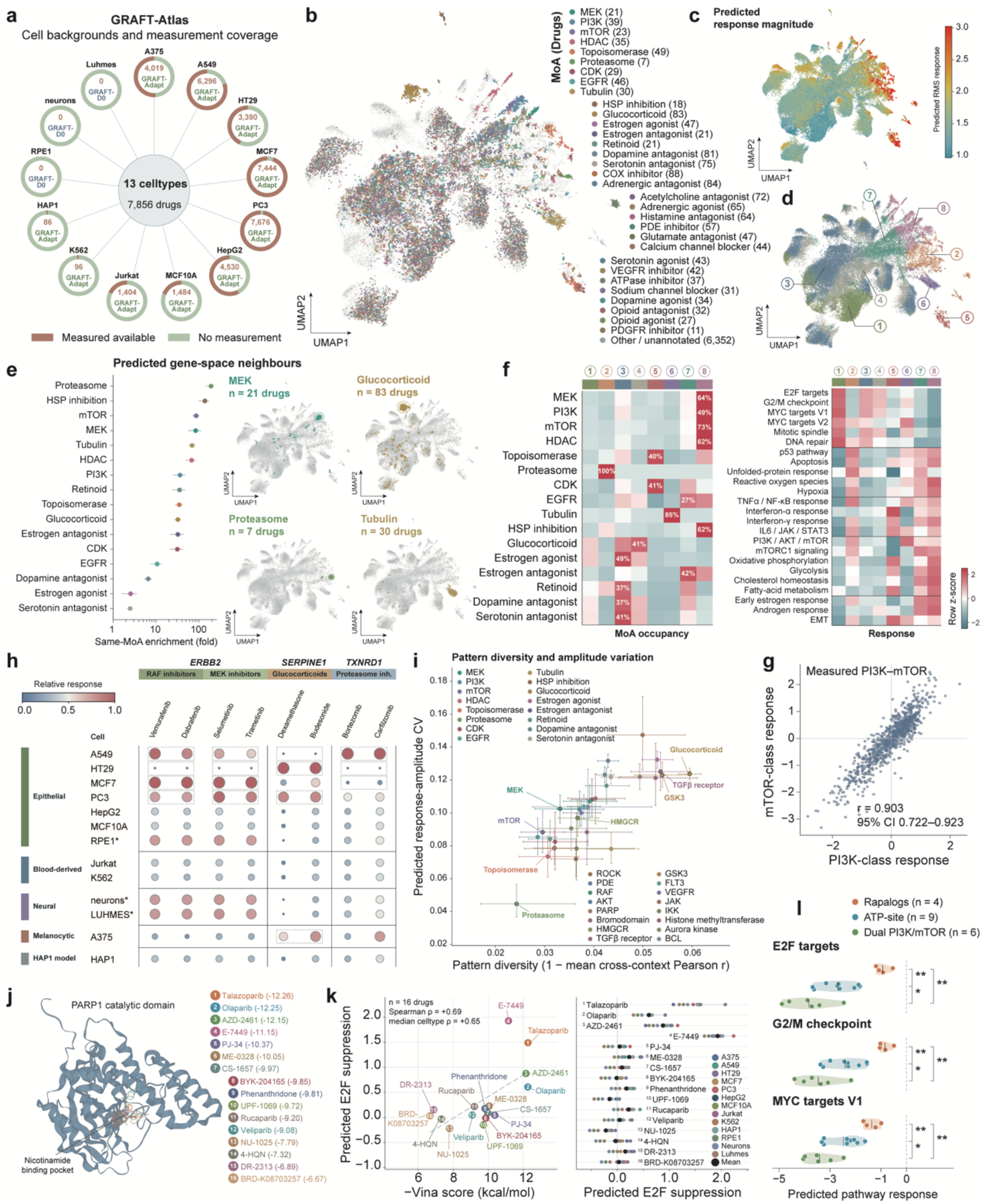
Pharmacological organization and context-dependent programs in GRAFT-Atlas. **a**, Measured-response coverage across 7,856 drug identifiers and 13 contexts. **b–d**, Prediction UMAP for 7,755 chemical entities, colored by mechanism of action (MoA; b), root-mean-square magnitude (RMS; c) or cluster (d); points represent drug–context pairs. **e**, Same-MoA enrichment among 30 neighbors in gene space (means, 95% drug-bootstrap intervals) and example density contours. **f**, Row-standardized MoA occupancy (numbers, maximum percentages) and Hallmark rank scores across eight clusters. **g**, Measured PI3K-only versus mTOR-only fingerprints, with Pearson correlation and drug-bootstrap 95% interval. **h**, Predicted *ERBB2, SERPINE1* and *TXNRD1* expression changes for eight drugs across 13 contexts. Color and bubble area encode within-column min–max scaling; blue indicates relatively smaller changes. Asterisks identify D0 contexts; other contexts use GRAFT-Adapt. **i**, Cross-context pattern diversity (1−mean Pearson *r*) versus RMS coefficient of variation; class means and 95% entity-bootstrap intervals. **j**,**k**, PARP1-pocket docking (j; PDB 4UND) and −Vina versus predicted E2F suppression (negated mean signed E2F score) for 16 drugs, with context-specific profiles (k). Scatter points average 13 contexts; horizontal ranges span three docking searches; *ρ*, Spearman correlation. **l**, Mean signed E2F, G2/M and MYC targets V1 predictions for rapalogs, ATP-site mTOR inhibitors and dual PI3K/mTOR inhibitors (*n* = 4, 9 and 6 drugs). Points are drug means across 13 contexts; violins show distributions and bars indicate subclass means. All pairwise comparisons use unadjusted two-sided exact Mann–Whitney tests: ^*^*P* < 0.05; ^**^*P* < 0.01.

Additional gains were associated with measured response heterogeneity. Here, gains denoted the query correlation difference between GRAFT-Adapt and GRAFT-D0. For drugs shared by the five cancer cell lines, we quantified heterogeneity as one minus the mean pairwise Pearson correlation of their complete 976-gene measured profiles. Higher values indicated less consistent responses across contexts. This measure correlated positively with mean adaptation gains (Spearman *ρ* = 0.455; **Fig. 4g**).

Drugs with greater measured cross-context heterogeneity therefore tended to benefit more from anchor-based adaptation, even after genetic conditioning.

Uniform manifold approximation and projection (UMAP) of measured responses showed that several drug–context pairs lay away from the main concentrations of their annotated pharmacological classes. The same embedding was colored by adaptation gain and measured response root mean square (RMS) (**Extended Data Fig. 5b**,**c**). These distinct patterns suggested context-dependent departures from typical class responses. Doxepin in Jurkat and tofacitinib in MCF10A showed stronger measured responses and larger adaptation gains than the same drugs in other cell lines (**Fig. 4h**). Amcinonide, orantinib and etoposide in Jurkat, and BRD-K68336408 in MCF10A, showed analogous class departures and stronger responses with larger gains (**Extended Data Fig. 5d**,**e**). Together, these cases showed that GRAFT-Adapt could use limited target drug measurements to improve prediction of distinctive responses in specific cellular contexts.

### GRAFT-Atlas organizes pharmacology and context-dependent responses

We constructed GRAFT-Atlas across 13 cellular contexts, covering 7,856 drug catalog identifiers and 102,128 drug–context combinations, including 65,703 without existing measurements. This expanded drug–context coverage 2.80-fold. Ten contexts used GRAFT-Adapt; RPE1, neurons and LUHMES used GRAFT-D0^5,21,22^. Prediction and measurement layers were kept separate and linked through common drug and context identifiers (**Fig. 5a**).

After merging identical chemical entities, we mapped predicted profiles for 7,755 entries across all 13 contexts, annotated by mechanism of action (MoA) or response magnitude (**Fig. 5b**,**c**). Clustering without MoA labels identified eight response groups (**Fig. 5d**). Same-MoA neighborhoods were also enriched in the full predicted gene space, with all drug-bootstrap interval lower bounds above the random enrichment ratio of 1 (**Fig. 5e**). Most MEK-, mTOR- and HDAC-inhibitor responses converged in cluster 8, together with approximately half the PI3K-inhibitor responses. Proteasome and microtubule inhibitors concentrated in clusters 2 and 6, respectively (**Fig. 5f**).

Hallmark scores based on within-profile gene ranks linked these groups to functional programs^23^. Cluster 8 had the lowest E2F, G2/M and MYC scores, consistent with shared proliferation suppression downstream of MEK–ERK and PI3K–mTOR signaling^17,24^. Cluster 2 combined low proliferation scores with unfolded-protein, TNFα/NF-κB and p53 responses (**Fig. 5f**). Additional pathway maps, class enrichment and gene fingerprints distinguished these stress programs (**Extended Data Fig. 6a–c**). Proteasome inhibitors had positive *HMOX1*/*SQSTM1* predictions, consistent with proteostasis stress^25^. HDAC inhibitors had positive *CDKN1A*/*GADD45A* predictions, consistent with growth arrest^26,27^. After excluding drugs annotated to both classes, PI3K and mTOR fingerprints remained correlated in the predicted (*r* = 0.966; **Extended Data Fig. 6h**) and measured (*r* = 0.903; **Fig. 5g**) layers.

**Figure 6.**
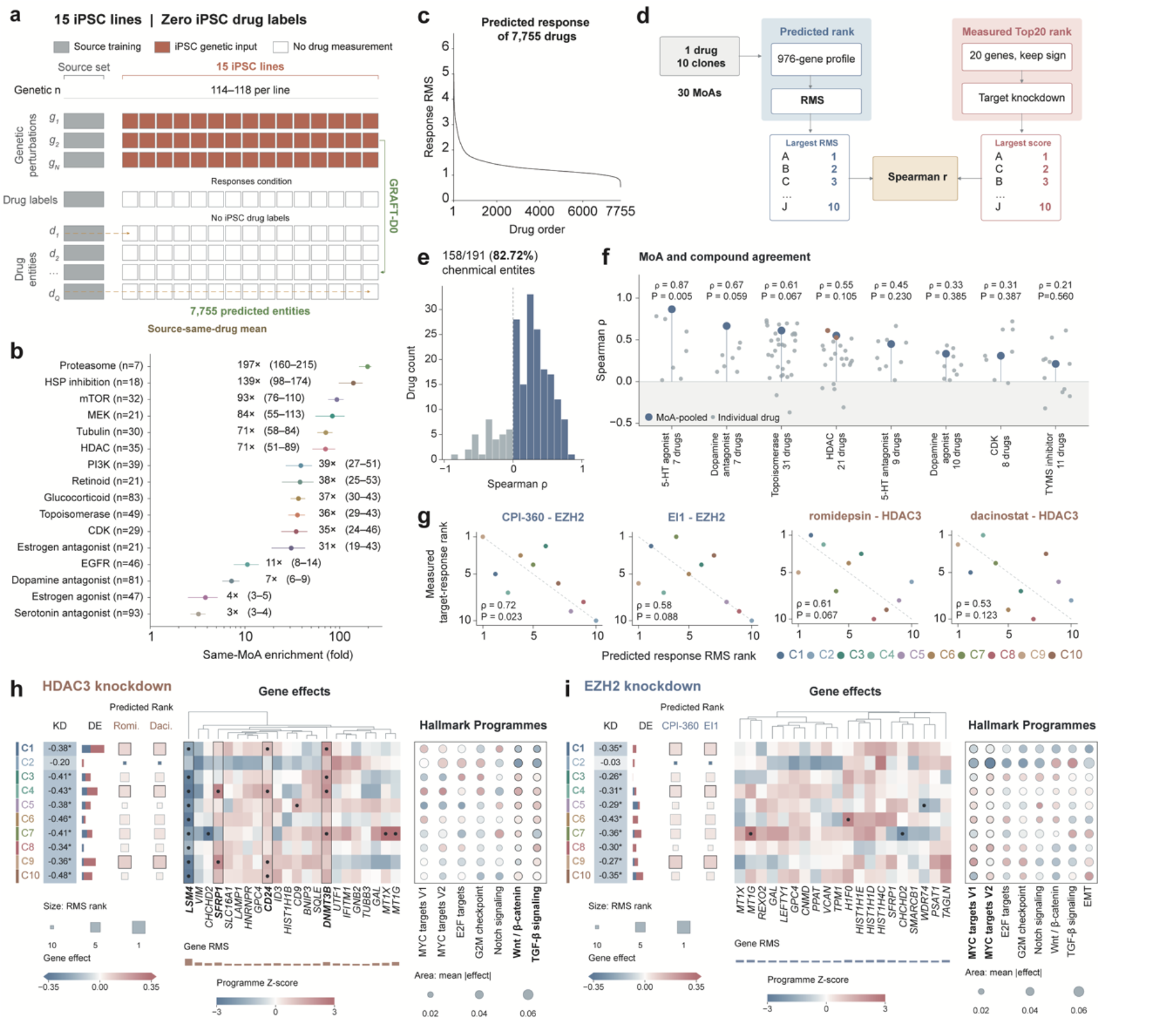
iPSC drug predictions and measured genetic-response correspondence. **a**, Prediction of 7,755 entities across 15 iPSC lines from ten donors using 114–118 genetic inputs per line, without iPSC drug measurements. **b**, Same-mechanism-of-action (MoA) enrichment among 30 neighbors; means and 95% entity-bootstrap confidence intervals (2,000 resamples). *n* denotes entities. **c**, Predicted root-mean-square (RMS) responses, ordered by their 15-line mean (black). d–i use ten separately selected iPSC lines, one from each donor. **d**, Predicted RMS ranks versus measured target-knockdown scores retaining source-drug Top20 directions. **e**, Distribution of donor-level correlations between predicted drug-response magnitudes and measured target-knockdown scores for 191 retained chemical entities. **f**, MoA-pooled (large dots) and member-drug (small dots) correlations across 9–10 donors. **g**, Four drug–target rank comparisons (*n* = 10 donors). **f**,**g**, Spearman *ρ* and exact two-sided nominal *P* values. **h**,**i**, Measured *HDAC3* and *EZH2* responses, respectively. Heatmaps show adjusted gene effects; dots and KD asterisks denote adjusted *q* < 0.05 as reported by the original study. DE bars count downregulated/upregulated genes (blue/red); lower bars show gene-effect RMS. Square size encodes predicted RMS rank (1, largest). Bubble color shows signed mean genetic effects standardized across donors; area shows unstandardized mean absolute effects. C1–C10 identify donors in g–i. Genetic comparisons reuse the model’s conditioning data.

We compared predicted changes in gene expression for eight drugs across 13 contexts (**Fig. 5h**). RAF/MEK inhibitors produced larger predicted increases in ERBB2 expression in A549, MCF7 and PC3 than in Jurkat and K562. Studies in BRAF-mutant thyroid cancer cells show that RAF/MEK inhibition can induce HER3 transcription and ERBB2/HER2 expression, promoting HER2–HER3 heterodimer signaling that bypasses MAPK-pathway blockade by activating parallel PI3K–AKT survival signaling. Dual EGFR/HER2 inhibition with lapatinib suppresses this rebound signal and enhances the growth-inhibitory effects of RAF or MEK inhibitors^28^. This provides a rationale for testing combined MAPK and HER2 inhibition in A549, MCF7 and PC3, where *ERBB2* induction was predicted to be stronger.

Glucocorticoids were predicted to induce stronger *SERPINE1* expression in HT29, PC3 and A375 than in the blood-derived lines Jurkat and K562, but weaker *SERPINE1* induction in A549. One possible explanation is that glucocorticoid receptor (GR) binding is shaped by chromatin accessibility. In mouse mammary 3134 and pituitary AtT-20 cells, GR mainly bound regions that were already open before hormone treatment. Because these cells had different pre-existing chromatin landscapes, GR occupied different genomic sites^12^. Differences in pre-existing chromatin accessibility may therefore influence which genes glucocorticoids activate and help explain the predicted variation in *SERPINE1* induction.

Under proteasome inhibition, predicted increases in TXNRD1 expression were greatest in A549, whereas HT29 and MCF7 showed smaller increases than Jurkat and K562. Because TXNRD1 is involved in antioxidant defense, its stronger predicted induction in A549 may reflect activation of a protective stress response. Related evidence from myeloma cells shows that glutathione buffering reduces bortezomib-induced stress responses and cytotoxicity, whereas xCT inhibition enhances bortezomib-induced cell death^25^. These findings support testing whether antioxidant-pathway blockade sensitizes A549 to proteasome inhibitors.

The atlas also separated cross-context variation in response pattern from variation in magnitude. GR agonists showed greater pattern diversity than mTOR inhibitors (**Fig. 5i**; **Extended Data Fig. 6d**). We docked 16 drugs with recorded PARP1 activity into the same nicotinamide-binding pocket using AutoDock Vina (**Fig. 5j**)^29^. More favorable scores corresponded to stronger predicted E2F suppression (Spearman *ρ* = 0.69), with the same direction across contexts (**Fig. 5k**). As docking scores were not model inputs, this association independently linked predicted transcriptional effects to structural rankings within a common target pocket.

All 41 GR-agonist groups had lower predicted ROS-associated transcriptional scores in A549 than their means across the other four cancer contexts (**Extended Data Fig. 6e**). The class-specific contrast remained negative after adjustment for non-GR responses (**Extended Data Fig. 6f**). GR-dependent repression of *SLC7A11* has been reported in A549 but not MCF7^30^. This suggests cell-specific antioxidant regulation, although the scores do not measure intracellular ROS levels.

The mTOR subclasses showed progressively more negative E2F, G2/M and MYC targets V1 predictions from rapalogs to ATP-site and dual PI3K/mTOR inhibitors (**Fig. 5l**), with structural references shown in **Extended Data Fig. 6g**. ATP-site inhibitors differed from rapalogs across all three programs after multiple-testing correction. This ordering is consistent with incomplete inhibition of 4E-BP1 phosphorylation by rapalogs and more complete inhibition of translational output by ATP-site inhibitors^31^. The further reduction with dual inhibitors suggests that broader inhibition along the PI3K–mTOR axis may deepen the downstream transcriptional response.

### GRAFT links iPSC drug-response predictions to measured genetic programs

We extended GRAFT to public genetic perturbation data from human induced pluripotent stem cell (iPSC) lines^32^. Without iPSC drug measurements, we predicted responses to 7,755 chemical entities across 15 lines from ten donors (**Fig. 6a**).

Same-MoA neighbor enrichment showed that predicted profiles retained pharmacological organization (**Fig. 6b**). Across these 15 lines, predicted response magnitudes showed broadly similar drug-level patterns when drugs were ordered by their mean response RMS (**Fig. 6c**).

For drugs whose molecular targets had corresponding Perturb-seq measurements, we examined whether donor rankings based on predicted drug-response magnitude agreed with rankings based on measured target-gene knockdown effects. For the EZH2 inhibitor CPI-360, we compared each donor’s predicted response RMS with the measured *EZH2*-knockdown score. Each genetic score summarized effects across the 20 genes with the strongest source responses to that drug, retaining source directions (**Fig. 6d**). A positive correlation indicated that donors with larger predicted drug responses tended to have higher genetic scores in the corresponding source-drug response direction. In the retained exploratory set, 158 of 191 entities (82.72%) showed positive correlations; all eight displayed MoA classes also had positive pooled associations (**Fig. 6e,f**). For example, predicted responses to CPI-360 and EI1 were paired with *EZH2*-knockdown effects^33,34^. Predictions for the HDAC inhibitors romidepsin and dacinostat were paired with *HDAC3*-knockdown effects^35,36^. Across-donor Spearman correlations for these four drugs ranged from 0.53 to 0.72 (**Fig. 6g**).

We then examined measured genes and programs in the Perturb-seq data used as model input. *HDAC3* and *EZH2* each showed significant on-target transcript reduction in nine lines. HDAC3 regulates hepatic differentiation of human pluripotent stem cells^37^. After its knockdown, *LSM4* decreased and *DNMT3B* increased in all ten lines, *CD24* increased in nine, and *SFRP1* varied in direction. C9, C1 and C4, the highest-ranked lines for romidepsin and dacinostat, showed significant *LSM4* reductions and *CD24* increases (author-adjusted *q* < 0.05; **Fig. 6h**). LSM4 participates in pre-mRNA processing, whereas CD24 is associated with the primed state of human pluripotent stem cells, providing interpretable downstream genetic readouts^38,39^.

*EZH2* knockdown produced more heterogeneous downstream effects. Its variable genes distinguished lines. C9 showed relatively high standardized MYC targets V1, E2F and G2/M scores alongside the highest predicted CPI-360 and EI1 responses (**Fig. 6i**). These findings suggest that GRAFT can identify candidate inter-donor differences in drug responses from measured genetic perturbation programs, providing a basis for further experimental evaluation in donor-derived iPSC.

## Discussion

GRAFT connects genetic-to-chemical transfer with adaptation across cellular contexts. A shared state derived from target genetic responses corrects source responses to multiple drugs without target drug measurements. When such measurements become available, GRAFT-Adapt transfers their residual errors to unmeasured queries. Internal and external evaluations support both stages, while genetic-budget analyses show that limited target measurements can provide informative context. Reusing a conditioning state across queries connects one target genetic screen to a broad panel of source-measured drugs. This approach expands the use of genetic screens from characterizing perturbation effects to predicting missing drug responses.

Genetic responses directly record functional changes following intervention, as demonstrated by large-scale Perturb-seq studies^5,20^. Genetic and chemical interventions can converge on shared downstream programs, providing a basis for transfer between response modalities^3,6^. Within a cellular context, both modalities act on interconnected regulatory networks. Their shared outputs allow differences in genetic responses to inform corrections for drug effects across contexts. Gains diminished when conditioning was disabled, target inputs were replaced with source-cell responses, or genetic residual correspondences were disrupted. These controls identify contextual information beyond source responses to the same drug. Most performance was retained in a compact state, without consistent benefits from greater architectural complexity. Conditioning on a perturbation set therefore provides context without requiring a genetic measurement matched to every query drug’s target.

GRAFT-Atlas extends comparisons of drug-induced transcriptional responses to previously unmeasured contexts, revealing shared and context-dependent patterns across pharmacological classes. Convergence on proliferation-associated outputs is consistent with MIX-Seq observations, but cannot by itself identify upstream mechanisms^1,40^. Differences in stress related programs and individual gene responses further distinguished pharmacological classes^41^. Drugs acting on different upstream targets can converge on common proliferation outputs while retaining class-specific stress and metabolic programs. For example, proteasome perturbation linked decreased proliferation-associated transcription to proteostasis stress. Distinct mTOR inhibitor groups further connected mode of action with the magnitude of downstream transcriptional changes. These differences generate testable hypotheses about how drugs with similar transcriptional effects may differ across cellular contexts. Genetic–chemical and combinatorial genetic perturbations, combined with longitudinal phenotypic readouts, could test whether these predicted relationships correspond to drug sensitivity or resistance^42-44^.

Transfer reliability depends on data quality, source coverage and comparability of experimental conditions^10,45^. HAP1 showed lower prediction correlations and no significant drug-anchor gains in paired tests. Its genetic inputs combined stimulation conditions, whereas drug signatures came from another platform, potentially weakening correspondence between modalities. Because evaluation metrics differ in their sensitivity to response patterns and magnitudes^46^, we considered correlations, magnitude errors and recovery of strongly responding genes together. Accumulating paired data under more consistent experimental conditions could help distinguish technical differences from biological context.

The iPSC analysis extends GRAFT to donor-derived cells, where comprehensive drug-response measurements are often unavailable. Predicted profiles retained same-MoA organization, and predicted drug-response magnitudes showed positive donor-level associations with selected measured target-knockdown effects. These findings suggest that an individual’s genetic response profile could help expand the drug-response map for that sample. Across-donor RMS rankings nevertheless largely reflected a shared tendency in response magnitude, and the functional comparisons reused genetic data supplied to the model. Independent drug-treatment experiments in separate iPSC samples are therefore needed to test predicted drug-specific differences. Broader application also depends on the contexts and response features represented during training. In its current transfer setting, GRAFT relies on cancer cell lines as source contexts and predicts mean transcriptional responses over a common gene panel. Incorporating iPSC or lineage-matched patient-derived source contexts could improve the reliability of comparisons between individuals. Extending coverage of dose, exposure duration and cellular states could further broaden the prediction scope, building on existing approaches to modeling cellular heterogeneity and response distributions^7,8^.

As perturbation datasets expand, GRAFT could be extended to investigate tissue- and individual-specific transcriptional responses associated with drug toxicity^8^. Prior studies have established transcriptomic liver-injury prediction, kidney organoid injury models and donor-dependent susceptibility to liver injury^47-49^. These applications motivate extending GRAFT to hepatocytes, kidney-related cells and their organoids to compare predicted stress, metabolic imbalance and injury-associated programs across tissues and individuals. Perturb-seq in primary human T cells further provides an experimental foundation for functional profiling of primary cells^50^. In these systems, responses to a limited gene panel could define a contextual representation, with a few drug measurements used for calibration. Integrating the resulting predictions with dose, exposure duration and measured toxicity phenotypes could connect transcriptional programs to cellular injury. Such a workflow could prioritize candidate drugs and individual differences in toxicity susceptibility for experimental testing while making more efficient use of limited patient material.

## Data availability

LINCS L1000 Level-5 signatures are available from CLUE (https://clue.io/data/CMap2020)^3^. GEO accession GSE264667 provides genetic perturbation data for HepG2 and Jurkat^18^. MCF10A data are available under GSE108699^19^. HAP1 data are available under GSE281048^20^. LUHMES data are available under GSE142078^22^. K562 and RPE1 data are available from Figshare at https://doi.org/10.25452/figshare.plus.20029387.v1^5^. The day-7 neuronal data from Tian et al. are available through scPerturb (https://doi.org/10.5281/zenodo.13350497)^21,45^. The iPSC count matrices and metadata (https://doi.org/10.6084/m9.figshare.27989294.v2) and covariate-adjusted genetic perturbation effects (https://doi.org/10.6084/m9.figshare.26819743.v1) were provided by Feng et al.^32^. Frozen perturbation representations were obtained from UniPert^6^. Hallmark gene sets (v2026.1.Hs) were obtained from MSigDB^23^. Structural coordinates are available from the Protein Data Bank (accessions 7M0X, 4JSX, 1FAP and 4UND), and PARP1 activity records from ChEMBL (target CHEMBL3105). Processed response matrices, atlas predictions and fixed analysis inputs are available in the GRAFT reproduction repository and its v0.1.0 release (https://github.com/Royoone/GRAFT-reproducibility/releases/tag/v0.1.0).

## Code availability

The GRAFT implementation, pretrained models and usage examples are available at https://github.com/Royoone/GRAFT. Code for reproducing the computational analyses and selected figures is available at https://github.com/Royoone/GRAFT-reproducibility.

## Author Contributions

C.L., M.H., and Q.R.L. conceived the study. C.L. and M.H. developed the methodology. C.L., M.H., Y.W. and X.L. curated the data. C.L. performed the formal analyses and developed the software. M.H., Y.W., D.G. and S.W. validated the results. C.L. wrote the original draft. X.L., M.H. and D.G. reviewed and edited the manuscript. X.L. and Q.R.L. supervised the study.

## Methods

### Prediction task and datasets

GRAFT predicts mean drug-induced transcriptional responses in a target cellular context from genetic perturbation responses measured in that context. Training additionally uses genetic and drug responses from source contexts. We distinguish three settings according to the availability of target-context drug measurements. In D0, these measurements are reserved for evaluation and excluded from training and model selection; source measurements of the same drugs remain available. GRAFT-Adapt uses measured responses to a disjoint set of drug anchors to calibrate query predictions. GRAFT-Atlas uses all available drug anchors to construct a response atlas.

### Source-context data

Source data comprised LINCS L1000 Level-5 response signatures for A375, A549, HT29, MCF7 and PC3^3,6^. Genetic perturbations were CRISPR loss-of-function perturbations profiled at 96 h, and drug treatments were profiled at 10 µM for 24 h. Responses were aligned to a fixed 976-gene LINCS panel, and signatures sharing a cellular context and perturbation identifier were averaged arithmetically. Only conditions with valid UniPert representations were retained. Level-5 signatures were used directly as signed responses without further control subtraction. The five contexts collectively covered 4,994 genetic targets and 7,856 drug identifiers, of which 4,156 targets and 3,220 drugs were measured in all five contexts.

### External-context data

External genetic perturbation data comprised HepG2 and Jurkat (GSE264667), K562 (Figshare 20029387), MCF10A (GSE108699) and HAP1 (GSE281048)^5,18–20^. For the first four contexts, raw counts were log1p-transformed, and mean non-targeting control expression was subtracted from mean expression for each genetic target. Perturbation groups required at least five cells; MCF10A analyses included only single-gene perturbations. HAP1 responses were calculated separately under IFNB, IFNG, INS, TGFB and TNFA stimulation: counts were averaged within perturbation groups before log1p transformation and control subtraction, and responses to each target were then averaged across stimulation conditions. Each dataset was aligned to the fixed gene panel, assigning zero to unmeasured genes.

External drug responses for evaluation were obtained from LINCS2020 beta Level-5 chemical signatures^3^. Within each context, we averaged available signatures for each drug across doses and time points. BRD identifiers, canonical names and aliases linked these drugs to the source catalog. The matched query sets contained 4,530 drugs in HepG2, 1,484 in MCF10A, 1,404 in Jurkat, 96 in K562 and 86 in HAP1.

Atlas construction additionally included RPE1 (Figshare 20029387), day-7 neurons from the Tian et al. dataset curated by scPerturb, and Luhmes (GSE142078)^5,21,22,45^. Counts were log1p-transformed, and mean control expression was subtracted from mean perturbed expression.

### Genetic conditioning of drug response predictions Perturbation representations and source references

Let *c* denote the target context, *b* a source context and *p* a genetic or drug perturbation. We reserve *d* for drugs and *g* for response genes. The identity embedding **x**_*p*_ ∈ ℝ^256^ and mean transcriptional response **y**_*cp*_ ∈ ℝ^*G*^ describe each perturbation, with *G* = 976. Non-bold symbols with subscript *g* denote individual gene components. Both genetic and drug identities are represented by frozen 256-dimensional UniPert embeddings^6^. The pretrained genetic encoder combines ESM2 protein sequence features with a graph neural network. The drug encoder projects 2,048-bit ECFP4 fingerprints through a fully connected layer.

The source reference summarizes the average response across available source contexts, relative to which GRAFT predicts a target-context correction. For perturbation *p*, ***µ***_*p*_ is the equally weighted mean of responses from source contexts measuring that perturbation. For genetic targets unmeasured in sources, we instead average the context-specific mean genetic responses.References and scaling use only source contexts allowed by the evaluation split; source-set subscripts are omitted below. The scale *s* is the square root of the mean squared drug response, first averaged within each source context and then equally across contexts. We use 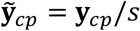 and 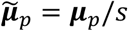 internally, and multiply final predictions by *s* to restore the original scale. Tildes denote responses in this scaled space throughout the model definition.

### Context representation from genetic response cross moments

To represent context-dependent responses, GRAFT relates perturbation identity to the deviation of its measured genetic response from the source reference. Consider a context *c* with *N*_*c*_ genetic support conditions, with *p*_*i*_ identifying the *i*-th perturbation. Its response residual is defined as 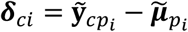, and the support-set mean residual as 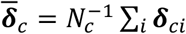. Identity and reference features are encoded separately from the mean-centered residual.

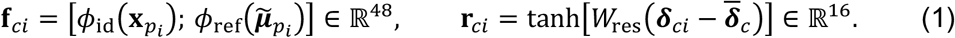

The semicolon denotes concatenation. The identity encoder *ϕ*_id_ applies layer normalization, a 256-to-32 linear projection and a tanh activation. The reference encoder *ϕ*_ref_ uses a 976-to-16 projection followed by tanh, while the residual projection *W*_ref_ ∈ ℝ^16×*G*^ is bias-free. Features are stacked across support conditions, centered columnwise and divided by their columnwise root mean square (RMS), with a lower bound of 0.05. This yields the support matrices 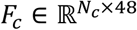 and 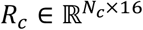.

Their cross-moment matrix summarizes the association between identity-and-reference features and genetic response residuals, independently of support ordering.

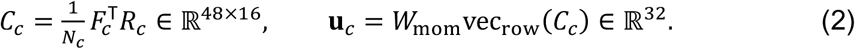

Here, vec_,67_ denotes row-major vectorization and *W*_565_ ∈ ℝ^32×768^ is a bias-free projection. We bound the RMS of this projection before mapping it to the context state.

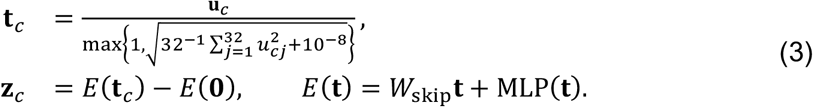

The state variables satisfy **t**_*c*_, **z**_*c*_ ∈ ℝ^32^; *u*_*cj*_ denotes component *j* of **u**_*c*_. The residual encoder combines a bias-free, identity-initialized 32-to-32 skip projection *W*_skip_ with a multilayer perceptron (MLP). The MLP comprises layer normalization, 32-to-64 and 64-to-32 linear layers, and an intervening GELU activation; its final weights and biases are initialized to zero. Without genetic support, **t**_*c*_ = **0**; a single support condition also gives zero cross-moments after centering. The state **z**_*c*_ is shared by all drug queries in the same context.

### Context dependent modulation and response prediction

GRAFT uses the genetic context state to modulate drug features through a hypernetwork and feature-wise linear modulation (FiLM). For drug *d*, concatenated identity **x**_*d*_ and source reference 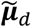 are encoded into 64-dimensional features **h**_*d*_ by a linear layer, GELU and layer normalization. The hypernetwork *H* comprises 32-to-32 and 32-to-128 linear layers separated by GELU. Its output defines two 64-dimensional vectors that scale and shift the drug features.

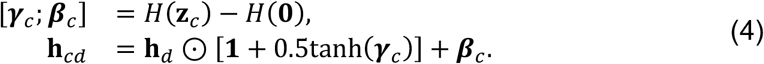

Here, ⊙ denotes elementwise multiplication and **1** the matching all-ones vector. The vectors ***γ***_*c*_ and ***β***_*c*_ determine scaling and shifting, respectively. With *ψ*(**h**) = GELU(LayerNorm(**h**)), prediction adds a context-specific correction to the source reference.

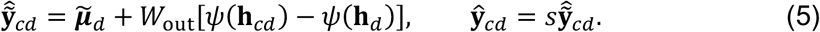

The bias-free readout is *W*_out_ ∈ ℝ^*G*×^64. Subtraction of the zero-state outputs and the difference-based readout ensure that predictions equal the source reference ***µ***_*d*_ when **z**_*c*_ = **0**. The backbone contains 216,752 trainable parameters.

An alternative, saturating state mapping was used for external-context generalization and analyses of genetic support budgets, source coverage and input substitution. This mapping takes the form 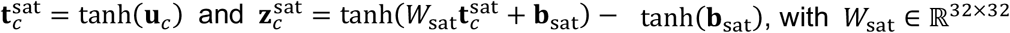 and 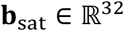, with *W*_sat_ ∈ ℝ^32×32^ and **b**_sat_ ∈ ℝ^32^. Genetic features, cross-moments and response readout otherwise follow equations (1)–(2) and (4)–(5). Each model was trained separately, including its own cross-moment projection producing **u**_*c*_. Internal genetic-budget and source-coverage experiments used 200 training steps.

### Model training

Training mimicked transfer to an unmeasured context through repeated source-context prediction tasks. For internal evaluation, each cancer cell line was held out in turn and the remaining four supplied training data. External predictions used all five cancer cell lines as sources. At each training step, one training context was randomly selected as a pseudo-target and the others supplied source references. All available genetic perturbations in the pseudo-target served as support, and 128 drugs were sampled as queries.

The training objective combines gene-weighted response error with agreement in response pattern. For *B* context–drug queries indexed by *q*, let 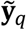 and 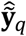 denote scaled measurements and predictions, respectively,

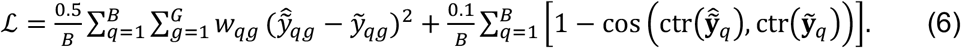

where ctr(**v**) = **v** − *G*^−1^<**1**^⊤^**v**=**1** centers a vector across genes and cos denotes cosine similarity. Query-specific gene weights satisfy *w* _*qg*_ ≥ 0 and ∑_*g*_ *w* _*qg*_ = 1.

Gene weights were derived from source signatures to distinguish each drug response from responses to other drugs in the same context. For each context–drug condition, the difference between the drug and remaining-drug means was divided by a shrinkage standard error. The shrinkage variance was the weighted mean of drug and reference variances, with respective weights max(*n*_*q*_ − 1,0) and 5. Here, *n*_*q*_ counts non-overlapping connected components defined by shared underlying experimental wells. The standard error was obtained by dividing the sum of shrinkage and reference variances by max(*n*_*q*_, 1) and then taking the square root. Reference variance was bounded below by 10^(0^. Absolute scores were min–max scaled across genes, squared and normalized to sum to one, giving *w* _*qg*_. Constant scores were assigned uniform weights.

We optimized GRAFT for 200 steps using AdamW with a learning rate of 0.001, weight decay of 10^−5^ and gradient-norm clipping at 1. Three independent random seeds were used, and final checkpoints were retained without early stopping.

### Drug anchor adaptation

GRAFT-Adapt calibrates D0 predictions using measured responses to drug anchors while keeping the backbone fixed. The conditioned features *ψ*(**h**_*cd*_) are L2-normalized for each drug and augmented with a constant, giving 65-dimensional features. For disjoint anchor and query sets *A* and *Q*, these form the row-wise matrices 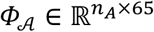 and 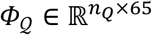, where *n*_S_ = |*A*| and *n*_*Q*_ = |*Q*|. Anchor residuals 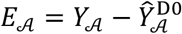 have dimension *n*_*A*_ × *G* and remain on the original response scale. Ridge regression transfers these residuals to query drugs.

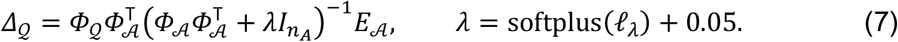

Here, 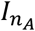 is the identity matrix, *l*_*λ*_ a learnable scalar and 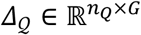 the predicted correction. Direct residual correction is combined with rank-8 gene mixing, parameterized by *U* ∈ ℝ^*G*×^8 and *V* ∈ ℝ^8×*G*^.

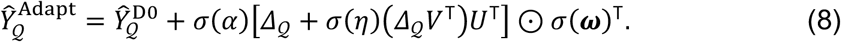

Here, *σ* denotes the logistic sigmoid. The learned scalars *α* and *η* control overall correction strength and low-rank mixing, respectively. Gene-specific gates ***ω*** ∈ ℝ^*G*^ are transposed and broadcast across queries. With no anchors, *Δ*_*Q*_ = 0, so GRAFT-Adapt returns the D0 prediction.

### Learning the adaptation parameters

Adaptation parameters were learned from anchor–query tasks within source contexts using frozen backbone features. Nonzero anchor budgets were sampled from 1, 2, 5, 10 and 20, and 15% of tasks contained no anchors. The objective combined Top20 and Top50 correlation losses, Top20 mean squared error (MSE), deviation from D0 across all genes, and a penalty for decreased correlation. Their respective weights were 1, 0.25, 0.05, 0.25 and 2. Zero-anchor tasks additionally included a penalty of 0.05*σ*(*α*).

Internal adapters used three non-target contexts for fitting and the fourth for checkpoint selection by Top20 correlation gain with 20 anchors. Training used Adam (learning rate 0.02, gradient clipping 5; 40 epochs, 16 tasks per epoch). Features came from ordinary leave-one-context-out backbones and were not regenerated for nested target splits. The external adapter used all five cancer cell lines. At inference, network weights were fixed and corrections were computed from available anchors using equation (7).

### Benchmark design and comparison methods

#### Query selection and D0 comparisons

All five folds and three random seeds shared a fixed query panel. The 3,220 drugs measured in all five cancer cell lines were ordered by a fixed SHA-256 rule based on drug identifiers; the first 20% (644 drugs) were used as queries, leaving 2,576 drugs for anchor sampling. D0 training and model selection excluded target drug responses while permitting source responses to the same drugs. External evaluation included all queries matching the source drug catalog. All methods used the same queries; drugs that could not be encoded received the source same-drug mean prediction.

Internal evaluation included 16 comparator configurations. Alongside the source same-drug mean, we evaluated task-adapted DeepCE, TranSiGen, PRnet, chemCPA and XPert, Morgan MLPs with 256- and 2,048-bit fingerprints, a UniPert MLP, and CIGER with hinge or RankNet ranking loss. G2CP was evaluated under D0, 20-anchor and 10% drug-budget settings, and HyperMap under D0 and 20-anchor settings. External evaluation included nine D0 comparators: the source same-drug mean, DeepCE, CIGER, TranSiGen, PRnet, chemCPA, Morgan MLP, XPert and G2CP.

Comparison networks were adapted to the available context information. DeepCE, CIGER, chemCPA and the MLP baselines used cell one-hot encodings, replaced by the mean source encoding for an unseen target. TranSiGen, PRnet and XPert did not use cell one-hot encodings. Where context expression was required, we supplied a constant mean source drug-response profile. For internal comparisons, this mean used only source-training rows. Networks were trained with Adam at a learning rate of 0.001, batch size of 256 and gradient-norm clipping at 1. CIGER’s gene-pair ranking loss and TranSiGen’s KL term had weights of 0.5 and 0.1, respectively.

G2CP used UniPert, 64-dimensional gene representations and a cross-gene readout. Source genetic and drug responses, target genetic responses and permitted non-query drug responses were pooled before random training–validation splitting, with each budget configuration trained independently from scratch. The 20-anchor configuration used the same anchors as HyperMap. The 10% budget was defined relative to all target drugs, with anchors sampled only from the non-query pool.

Target query responses were excluded from training and model selection throughout. HyperMap used a LINCS adaptation of the authors’ network, with a zero control vector and drug one-hot encodings, and was trained by second-order model-agnostic meta-learning (MAML) for 50 outer epochs. Adaptation with 20 anchors used three full-network updates, with backtracking to reduce the step size according to anchor loss, without using query responses.

#### Anchor budgets and shared mechanisms of action

Internally, anchors were drawn from 2,576 non-query drugs. External data were split into fixed query and candidate pools of approximately 80% and 20%, with prespecified examples retained as queries. All ten contexts used 0, 1, 2, 5, 10 and 15 anchors, with 10 as the primary budget and higher budgets where candidates permitted. Thirty random drug orderings per context defined nested anchor sets shared across the three backbone seeds and all adaptation methods.

GRAFT-Adapt was compared with standard and gated ridge regression. Internally, we also tested a gated kernel using UniPert cosine similarity to weight anchor residuals. Queries were grouped by shared, unshared or missing MoA annotations relative to each anchor list. Rankings equally weighted Pearson correlation, MSE and MAE over Top20 and Top50 windows. These six metrics were selected post hoc from the LINCS comparison; all eighteen metrics are provided in the source analysis tables.

#### Model components and information requirements Component ablation and alternative architectures

With weights and queries fixed, we separately disabled FiLM scaling, shifting or the context state, retained state dimensions selected by magnitude or at random while setting the remainder to zero, and performed response permutation and identity-matched source substitution. Raw responses were permuted before reference subtraction, whereas residual permutation preserved identity–reference pairs.

Alternative architectures were retrained under the same five folds, three seeds and 200-step budget, varying state and readout widths, hypernetwork depth, cross-moment dimensionality, set encoding, gene graphs, drug–genetic cross-attention, low-rank readouts, alternative inputs and repeated modulation. Gene graphs were constructed only from training-source responses.

#### Genetic support and source coverage

Genetic-budget experiments kept the backbone fixed and used nested random support subsets ranging from zero to all available conditions. Budget curves report means and standard deviations across three seeds after averaging subsamples within each seed. Source coverage compared all two-source and three-source combinations with four-source models. Correct-input and source-substitution tests used matched conditions among the 4,156 shared genetic targets, whereas shuffled-input tests used the complete target-context genetic set. Coverage and substitution scores were first averaged within each target context and then summarized as means and standard deviations across the five contexts.

#### Representation analyses

Ridge probes assessed how well alternative features predicted target-context drug residuals relative to the source same-drug mean. Features were row-L2-normalized and augmented with a constant, with ridge regularization fixed at 1. Disjoint fitting/query sets contained 2,576/644 drugs for full-coverage features and 1,350/364 drugs for mean drug-target protein representations. Identity substitution assessed compatibility with UniPert space independently of response labels. We fitted an ordinary least-squares map with an intercept from non-query identity embeddings to UniPert embeddings **(Extended Data Fig. 2f)**.

### Prediction accuracy and response structure

#### Evaluation metrics

Prediction accuracy was evaluated on mean drug–context response profiles. Strong-response windows selected genes with the largest absolute measured responses in the target context, with window sizes satisfying *K* ∈ {20,50,100}. Pearson and Spearman correlations, MSE and MAE were calculated on signed responses within each window. TopK recall was the overlap between measured and predicted absolute-response TopK gene sets divided by *K*. Positive and negative precision similarly used genes with the largest and smallest signed responses, respectively. Sign20 measured sign agreement within the measured Top20 window, while all-gene correlation used the complete gene panel (*G* = 976). Response magnitude was summarized by root mean square (RMS),

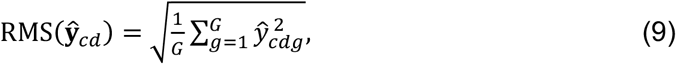

with the same definition applied to measured responses. Evaluation windows were defined separately from training weights, iPSC source-directed scoring windows and genetic-effect heatmap gene sets.

Internal Top20 correlations used the 641 queries with non-constant measured windows in all five contexts; other metrics retained all 644 queries. Constant measured windows produced missing correlations, whereas constant predictions against non-constant measurements were assigned zero correlation. Unless predictions were averaged before scoring, metrics were averaged across drugs within each context and seed, then equally across seeds and contexts.

Benchmarks, extended-window and drug-budget comparisons scored each seed separately before averaging, as did drug-level associations between source-mean performance and GRAFT gains. MoA, magnitude and signature-support stratification, discovery–confirmation comparisons and representative-drug analyses instead scored predictions averaged across three seeds. Internal and external method rankings equally averaged ranks across 21 and twelve metrics, respectively. Anchor-adaptation rankings used the six metrics specified above.

#### Response structure and prediction gains

For perturbations shared across the five cancer cell lines, cross-context correlations were calculated within each starting context’s absolute-response TopK genes and averaged first across ordered context pairs and then across perturbations. Total sums of squares were partitioned into a shared component, based on deviations of perturbation-specific cross-context means from the grand mean and weighted by the number of contexts, and a component capturing deviations of individual contexts from each perturbation mean.

Identity embeddings for 527 drugs in 27 classes and 524 genetic targets in six classes were clustered using KMeans. Agreement with annotations was evaluated by normalized mutual information (NMI) and adjusted Rand index (ARI) across 20 random seeds. Continuous features were row-L2-normalized, whereas ECFP4 remained binary. Genetic–drug correspondence was scored on genetic-response Top20 genes.

Response pattern difference was defined as one minus the mean pairwise full-profile Pearson correlation for the same drug across contexts. Together with RMS, it was used to analyze prediction gains of GRAFT over source means and of GRAFT-Adapt over D0.

Response maps used row-centered, L2-normalized profiles, followed by 50-component principal component analysis (PCA) and UMAP; annotations were excluded from fitting. For representative drugs, performance gains over equally weighted same-drug means from other contexts were assessed by paired *t*-tests across three model seeds. Measured magnitudes were calibrated against 56 shared drugs. Empirical tail probabilities were calculated using other drugs from the same contexts, with drugs rather than genes as replicates.

#### Response atlas and pharmacological analyses

GRAFT-Atlas extends prediction to 7,856 drug identifiers across thirteen cellular contexts. We averaged predictions across three seeds, using four-source backbones for the five cancer cell lines and five-source backbones elsewhere. Ten contexts used all available measured drugs as GRAFT-Adapt anchors; RPE1, neurons and Luhmes used D0 predictions. Full-anchor calibration used the equivalent 65-dimensional primal ridge solution. This resource-generation setting was separate from held-out query evaluation.

Predicted and measured responses were retained as separate layers, with unmeasured entries left missing in the measured layer. Entries sharing a complete InChIKey were averaged within each context; entries without valid keys retained their original identifiers. This produced 7,755 chemical entities. Maps, partitions and functional analyses used predicted responses, and class analyses retained overlapping MoA annotations.

To map pharmacological organization, we row-centered and L2-normalized predictions before randomized PCA with 50 components. UMAP used 200 neighbors and minimum distance 0.50, and MiniBatchKMeans partitioned PCA space into eight regions. MoA labels were excluded from fitting. Pharmacological enrichment was evaluated using 30 distinct neighbors per entity in the full 976-gene space, excluding self-matches. We averaged same-MoA neighbor fractions equally across contexts and entities. Fold enrichment was this average divided by the fraction of same-MoA entities among all other entities.

#### Functional programs and context specificity

Analyses used 49 MSigDB Hallmark (v2026.1.Hs) sets with at least five model genes^23^. Atlas scores averaged 2(*r* − 0.5)/*G* − 1 within each set; *r* denotes the ascending rank of signed responses, with average ranks for ties. Class fingerprints averaged raw signed responses within contexts and then across equally weighted contexts. Standardization was for heatmap display only. Fingerprint correlations excluded shared MoA members; measured PI3K-only/mTOR-only comparisons used entities measured in all five cancer lines.

One-sided hypergeometric tests evaluated the highest and lowest 50 genes of each class fingerprint against the 976-gene background. BH correction jointly covered all 21,952 tests. Context variability was measured across thirteen contexts by response pattern difference and the RMS coefficient of variation (standard deviation/mean), then summarized by MoA.

Drug-level mean signed program scores across thirteen contexts were compared by two-sided exact Mann–Whitney tests. ATP-site-versus-rapalog comparisons were Holm-adjusted across seven programs; Fig. 5l shows the unadjusted pairwise *P* values.

Raw signed gene-set scores were averaged within InChIKey connectivity groups, retaining all MoA annotations. Each group’s context-specific score was expressed relative to its mean in the remaining contexts. Welch’s t-tests compared these deviations between class members and non-members. BH correction covered 153,664 tests across the thirteen-context, ten-context GRAFT-Adapt and five-cancer-line analyses. Context-averaged comparisons across 112 classes and 49 programs were corrected separately.

#### Structural comparisons and molecular docking

Structural references used PDB 7M0X (PD-0325901), 4JSX (mTOR ATP site) and 1FAP (FKBP12–rapamycin–FRB). Rapalogs were illustrated by shared-core alignment to the sirolimus co-crystal; ATP-site and dual inhibitors were docked into 4JSX. One recorded three-dimensional configuration was used when ring configuration was unspecified.

PARP1 analysis included 16 atlas drugs with at least two ChEMBL CHEMBL3105 records for at least one measurement type (*K*_*d*_, *K*_*i*_ or IC_50_). Activities were summarized by the median for each type. Ligands were prepared with ETKDGv3/MMFF and Meeko 0.8.0, and docked into the fixed 4UND nicotinamide pocket using Vina 1.2.7. The search box was centered on the co-crystal ligand’s heavy-atom centroid, adding 12 Å to its dimensions (minimum 20 Å per axis).

Searches used exhaustiveness 32, up to ten poses and three seeds. Redocking required a top-pose symmetry-corrected heavy-atom RMSD below 2 Å.

We correlated negative mean top-pose Vina scores with E2F suppression (negated signed E2F score) using Spearman correlations. E2F scores were averaged equally across thirteen contexts and analyzed separately within each context. Analyses also adjusted for heavy-atom count by linear residualization. Ranges across three searches summarized computational variability. Docking provided independent structural references and was not used for training or prediction.

### iPSC drug predictions and genetic response correspondence

#### iPSC data processing and prediction

Day-3 CRISPRi data from Feng et al. covered nineteen iPSC lines, ten healthy donors and 444 targets^32^. We used published count matrices, metadata and covariate-adjusted effects with adjusted *P* values. Metadata-matched cells required positive UMI counts and at least 100 detected genes. Counts were merged by gene symbol, normalized to 10,000 UMIs per cell and log1p-transformed. Target responses were mean expression differences from same-line NonTarget controls, excluding guide-unassigned cells. All 976 model genes were measured.

Of these targets, 118 matched the source catalog and UniPert embeddings. Inputs required at least 20 targeted cells, two guides, 50 same-line controls and complete finite responses. Five-source GRAFT predictions used all eligible inputs and were averaged across three random seeds, without iPSC drug training or adaptation.

Excluding four quality-flagged lines left fifteen lines from ten donors with 114–118 inputs per line for pharmacological organization. MoA enrichment used the 30-neighbor definition with equal line/entity weights. PCA/t-SNE fitting excluded line and MoA labels; RMS used the original scale.

#### Source directed genetic scores

We explored whether variation in predicted drug-response magnitude across iPSC lines corresponded to variation in measured responses to matched genetic perturbations. This analysis used a separately defined set of ten lines, one per donor. Lines, drug predictions and genetic-score directions were fixed for subsequent analyses. Genetic scoring required at least ten targeted cells, two guides and 50 same-line controls. Let *ℐ*_*d*_ denote the absolute-response Top20 genes of source reference ***µ***_*d*_, and *T*_*cd*_ the valid targets matching drug *d* in line *c*. The value *e*_*ctg*_ is the control-referenced normalized response of gene *g* after perturbing target *t* in line *c*. We defined the source-directed genetic score as

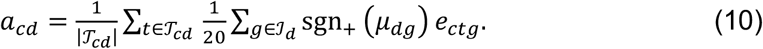

Here, sgn_B_ equals +1 for non-negative values and −1 otherwise. Targets were equally weighted, and scores were missing when no eligible target was available. Gene windows and directions were determined from sources, without selection using target-line correlations. These count-derived responses were distinct from the published covariate-adjusted effects.

#### Drug selection and association testing

Associations required paired RMS and genetic scores in at least five lines, yielding 235 evaluable entities across eight to ten lines. Spearman correlations were calculated across available lines. Association summaries retained 191 entities after excluding seven with unsuitable drug–target pairings and 37 lacking target-knockdown support. Knockdown support required a negative target-gene effect with published adjusted *P* < 0.05 for any matched target in any selected line meeting the scoring thresholds. All 235 entities and their filtering flags were retained in source data.

#### Statistical analysis and software

Statistical analyses used perturbations, drugs, chemical entities or connectivity groups as sampling units, retaining each unit’s records across contexts together. Model and docking seeds represented computational replicates. Random procedures used fixed seeds recorded in the analysis scripts. Bootstrap intervals were defined by the 2.5th and 97.5th percentiles. Resample counts were 2,000 for drug, fingerprint and iPSC enrichment analyses; 4,000 for adaptation gains; 5,000 for class variability and connectivity groups; and 10,000 for mTOR subclasses.

Adaptation comparisons averaged model seeds and anchor lists within each drug before applying two-sided paired Wilcoxon tests. Confidence intervals jointly resampled shared drugs for internal analyses and sampled drugs within contexts for external analyses. Holm correction covered ten context comparisons or thirty MoA-stratified comparisons. Anchor-budget shading shows the empirical 95% range across thirty anchor-list means, after averaging three seeds within each list, rather than a bootstrap interval.

All tests were two-sided except hypergeometric enrichment tests. Joint Holm correction covered the four internal magnitude/discovery–confirmation tests. BH correction was applied separately to 4,757 eligible drug–context case comparisons and 1,792 atlas variability tests. The latter spanned 112 classes, four context scopes and four metrics: pattern difference, RMS coefficient of variation, mean RMS and standard deviation of relative log2 RMS. Other multiple-testing families are defined alongside their corresponding analyses above.

GRAFT-Atlas analyses used Python 3.12.3, PyTorch 2.5.1, NumPy 1.26.4, pandas 2.2.3, SciPy 1.14.1, scikit-learn 1.5.2 and umap-learn 0.5.7. Additional analyses used openTSNE for dimensionality reduction and RDKit for chemical processing.

## References

1 McFarland, J. M. et al. Multiplexed single-cell transcriptional response profiling to define cancer vulnerabilities and therapeutic mechanism of action. Nat Commun 11, 4296 (2020). 10.1038/s41467-020-17440-w

2 Zhang, J. et al. Tahoe-100M: Mapping drug-induced molecular phenotypes at single-cell resolution. Cell 189, 5945–5961.e5948 (2026). 10.1016/j.cell.2026.08.035

3 Subramanian, A. et al. A Next Generation Connectivity Map: L1000 Platform and the First 1,000,000 Profiles. Cell 171, 1437–1452.e1417 (2017). 10.1016/j.cell.2017.10.049

4 Hetzel, L., Boehm, S., Kilbertus, N., Günnemann, S. & Theis, F. Predicting cellular responses to novel drug perturbations at a single-cell resolution. Advances in Neural Information Processing Systems 35, 26711–26722 (2022).

5 Replogle, J. M. et al. Mapping information-rich genotype-phenotype landscapes with genome-scale Perturb-seq. Cell 185, 2559–2575.e2528 (2022). 10.1016/j.cell.2022.05.013

6 Li, Y. et al. UniPert-G2CP bridges genetic and chemical screens from molecular representation to phenotype modeling. Cell 189, 4946–4963.e4948 (2026). 10.1016/j.cell.2026.06.005

7 Guo, Y. et al. Modelling drug-induced cellular perturbation responses with a biologically informed dual-branch transformer. Nature Machine Intelligence 8, 96–112 (2026). 10.1038/s42256-025-01165-w

8 Adduri, A. K. et al. Predicting cellular responses to perturbation across diverse contexts with State. Cell 189, 5914–5931.e5920 (2026). 10.1016/j.cell.2026.07.052

9 Dhaka, B., Gao, J. & Ideker, T. HyperMap: An Efficient Framework for Transferring Perturbation Responses Across Diverse Biological Contexts. bioRxiv, 2026.2004.2023.720505 (2026). 10.64898/2026.04.23.720505

10 Wei, Z. et al. Benchmarking algorithms for generalizable single-cell perturbation response prediction. Nat Methods 23, 451–464 (2026). 10.1038/s41592-025-02980-0

11 Perez, E., Strub, F., De Vries, H., Dumoulin, V. & Courville, A. in Proceedings of the AAAI conference on artificial intelligence.

12 John, S. et al. Chromatin accessibility pre-determines glucocorticoid receptor binding patterns. Nat Genet 43, 264–268 (2011). 10.1038/ng.759

13 Qi, X. et al. Predicting transcriptional responses to novel chemical perturbations using deep generative model for drug discovery. Nat Commun 15, 9256 (2024). 10.1038/s41467-024-53457-1

14 Pham, T. H., Qiu, Y., Zeng, J., Xie, L. & Zhang, P. A deep learning framework for high-throughput mechanism-driven phenotype compound screening and lits application to COVID-19 drug repurposing. Nat Mach Intell 3, 247–257 (2021). 10.1038/s42256-020-00285-9

15 Pham, T. H. et al. Chemical-induced gene expression ranking and its application to pancreatic cancer drug repurposing. Patterns (N Y) 3, 100441 (2022). 10.1016/j.patter.2022.100441

16 Tong, X. et al. Deep representation learning of chemical-induced transcriptional profile for phenotype-based drug discovery. Nat Commun 15, 5378 (2024). 10.1038/s41467-024-49620-3

17 Pratilas, C. A. et al. (V600E)BRAF is associated with disabled feedback inhibition of RAF-MEK signaling and elevated transcriptional output of the pathway. Proc Natl Acad Sci U S A 106, 4519–4524 (2009). 10.1073/pnas.0900780106

18 Nadig, A. et al. Transcriptome-wide analysis of differential expression in perturbation atlases. Nat Genet 57, 1228–1237 (2025). 10.1038/s41588-025-02169-3

19 Hill, A. J. et al. On the design of CRISPR-based single-cell molecular screens. Nat Methods 15, 271–274 (2018). 10.1038/nmeth.4604

20 Jiang, L. et al. Systematic reconstruction of molecular pathway signatures using scalable single-cell perturbation screens. Nat Cell Biol 27, 505–517 (2025). 10.1038/s41556-025-01622-z

21 Tian, R. et al. CRISPR Interference-Based Platform for Multimodal Genetic Screens in Human iPSC-Derived Neurons. Neuron 104, 239–255.e212 (2019). 10.1016/j.neuron.2019.07.014

22 Lalli, M. A., Avey, D., Dougherty, J. D., Milbrandt, J. & Mitra, R. D. High-throughput single-cell functional elucidation of neurodevelopmental disease-associated genes reveals convergent mechanisms altering neuronal differentiation. Genome Res 30, 1317–1331 (2020). 10.1101/gr.262295.120

23 Liberzon, A. et al. The Molecular Signatures Database (MSigDB) hallmark gene set collection. Cell Syst 1, 417–425 (2015). 10.1016/j.cels.2015.12.004

24 Elkabets, M. et al. mTORC1 inhibition is required for sensitivity to PI3K p110α inhibitors in PIK3CA-mutant breast cancer. Sci Transl Med 5, 196ra199 (2013). 10.1126/scitranslmed.3005747

25 Starheim, K. K. et al. Intracellular glutathione determines bortezomib cytotoxicity in multiple myeloma cells. Blood Cancer J 6, e446 (2016). 10.1038/bcj.2016.56

26 Richon, V. M., Sandhoff, T. W., Rifkind, R. A. & Marks, P. A. Histone deacetylase inhibitor selectively induces p21WAF1 expression and gene-associated histone acetylation. Proc Natl Acad Sci U S A 97, 10014–10019 (2000). 10.1073/pnas.180316197

27 Chen, Z. et al. Induction and superinduction of growth arrest and DNA damage gene 45 (GADD45) alpha and beta messenger RNAs by histone deacetylase inhibitors trichostatin A (TSA) and butyrate in SW620 human colon carcinoma cells. Cancer Lett 188, 127–140 (2002). 10.1016/s0304-3835(02)00322-1

28 Montero-Conde, C. et al. Relief of feedback inhibition of HER3 transcription by RAF and MEK inhibitors attenuates their antitumor effects in BRAF-mutant thyroid carcinomas. Cancer Discov 3, 520–533 (2013). 10.1158/2159-8290.Cd-12-0531

29 Eberhardt, J., Santos-Martins, D., Tillack, A. F. & Forli, S. AutoDock Vina 1.2.0: New Docking Methods, Expanded Force Field, and Python Bindings. J Chem Inf Model 61, 3891–3898 (2021). 10.1021/acs.jcim.1c00203

30 Lacher, S. E. et al. The Glucocorticoid Receptor Inhibits NRF2-Mediated Expression of SLC7A11. Free Radic Biol Med 241, 53–63 (2025). 10.1016/j.freeradbiomed.2025.09.008

31 Thoreen, C. C. et al. An ATP-competitive mammalian target of rapamycin inhibitor reveals rapamycin-resistant functions of mTORC1. J Biol Chem 284, 8023–8032 (2009). 10.1074/jbc.M900301200

32 Feng, C. et al. A genome-scale single-cell CRISPRi map of trans gene regulation across human pluripotent stem cell lines. Cell Genom 6, 101076 (2026). 10.1016/j.xgen.2025.101076

33 Bradley, W. D. et al. EZH2 inhibitor efficacy in non-Hodgkin’s lymphoma does not require suppression of H3K27 monomethylation. Chem Biol 21, 1463–1475 (2014). 10.1016/j.chembiol.2014.09.017

34 Qi, W. et al. Selective inhibition of Ezh2 by a small molecule inhibitor blocks tumor cells proliferation. Proc Natl Acad Sci U S A 109, 21360–21365 (2012). 10.1073/pnas.1210371110

35 Bradner, J. E. et al. Chemical phylogenetics of histone deacetylases. Nat Chem Biol 6, 238–243 (2010). 10.1038/nchembio.313

36 Khan, N. et al. Determination of the class and isoform selectivity of small-molecule histone deacetylase inhibitors. Biochem J 409, 581–589 (2008). 10.1042/bj20070779

37 Li, S. et al. Genetic and Chemical Screenings Identify HDAC3 as a Key Regulator in Hepatic Differentiation of Human Pluripotent Stem Cells. Stem Cell Reports 11, 22–31 (2018). 10.1016/j.stemcr.2018.05.001

38 Achsel, T. et al. A doughnut-shaped heteromer of human Sm-like proteins binds to the 3’-end of U6 snRNA, thereby facilitating U4/U6 duplex formation in vitro. Embo j 18, 5789–5802 (1999). 10.1093/emboj/18.20.5789

39 Shakiba, N. et al. CD24 tracks divergent pluripotent states in mouse and human cells. Nat Commun 6, 7329 (2015). 10.1038/ncomms8329

40 Szalai, B. et al. Signatures of cell death and proliferation in perturbation transcriptomics data-from confounding factor to effective prediction. Nucleic Acids Res 47, 10010–10026 (2019). 10.1093/nar/gkz805

41 Schubert, M. et al. Perturbation-response genes reveal signaling footprints in cancer gene expression. Nat Commun 9, 20 (2018). 10.1038/s41467-017-02391-6

42 McFaline-Figueroa, J. L. et al. Multiplex single-cell chemical genomics reveals the kinase dependence of the response to targeted therapy. Cell Genom 4, 100487 (2024). 10.1016/j.xgen.2023.100487

43 Norman, T. M. et al. Exploring genetic interaction manifolds constructed from rich single-cell phenotypes. Science 365, 786–793 (2019). 10.1126/science.aax4438

44 Coelho, M. A. et al. Base editing screens define the genetic landscape of cancer drug resistance mechanisms. Nat Genet 56, 2479–2492 (2024). 10.1038/s41588-024-01948-8

45 Peidli, S. et al. scPerturb: harmonized single-cell perturbation data. Nat Methods 21, 531–540 (2024). 10.1038/s41592-023-02144-y

46 Ahlmann-Eltze, C., Huber, W. & Anders, S. Deep-learning-based gene perturbation effect prediction does not yet outperform simple linear baselines. Nat Methods 22, 1657–1661 (2025). 10.1038/s41592-025-02772-6

47 Kohonen, P. et al. A transcriptomics data-driven gene space accurately predicts liver cytopathology and drug-induced liver injury. Nat Commun 8, 15932 (2017). 10.1038/ncomms15932

48 Vanslambrouck, J. M. et al. Enhanced metanephric specification to functional proximal tubule enables toxicity screening and infectious disease modelling in kidney organoids. Nat Commun 13, 5943 (2022). 10.1038/s41467-022-33623-z

49 Koido, M. et al. Polygenic architecture informs potential vulnerability to drug-induced liver injury. Nat Med 26, 1541–1548 (2020). 10.1038/s41591-020-1023-0

50 Schmidt, R. et al. CRISPR activation and interference screens decode stimulation responses in primary human T cells. Science 375, eabj4008 (2022). 10.1126/science.abj4008

