## Extended Data Fig. 1-6 for "Cross-modal transfer enables zero-shot prediction of drug responses across cell types"

† Drs. Luo and Hu contribute equally.

Qing Rex Lyu

### Contents

**Extended Data Fig. 1** Cross-context response conservation and genetic–chemical relationships.

**Extended Data Fig. 2** Genetic conditioning, capacity controls and representation probes.

**Extended Data Fig. 3** Detailed LINCS prediction benchmarks and pharmacological examples.

**Extended Data Fig. 4** Extended external benchmarks and genetic-input requirements.

**Extended Data Fig. 5** Adaptation gains and context-dependent measured drug responses.

**Extended Data Fig. 6** Predicted functional programs, context effects and mTOR structural references.

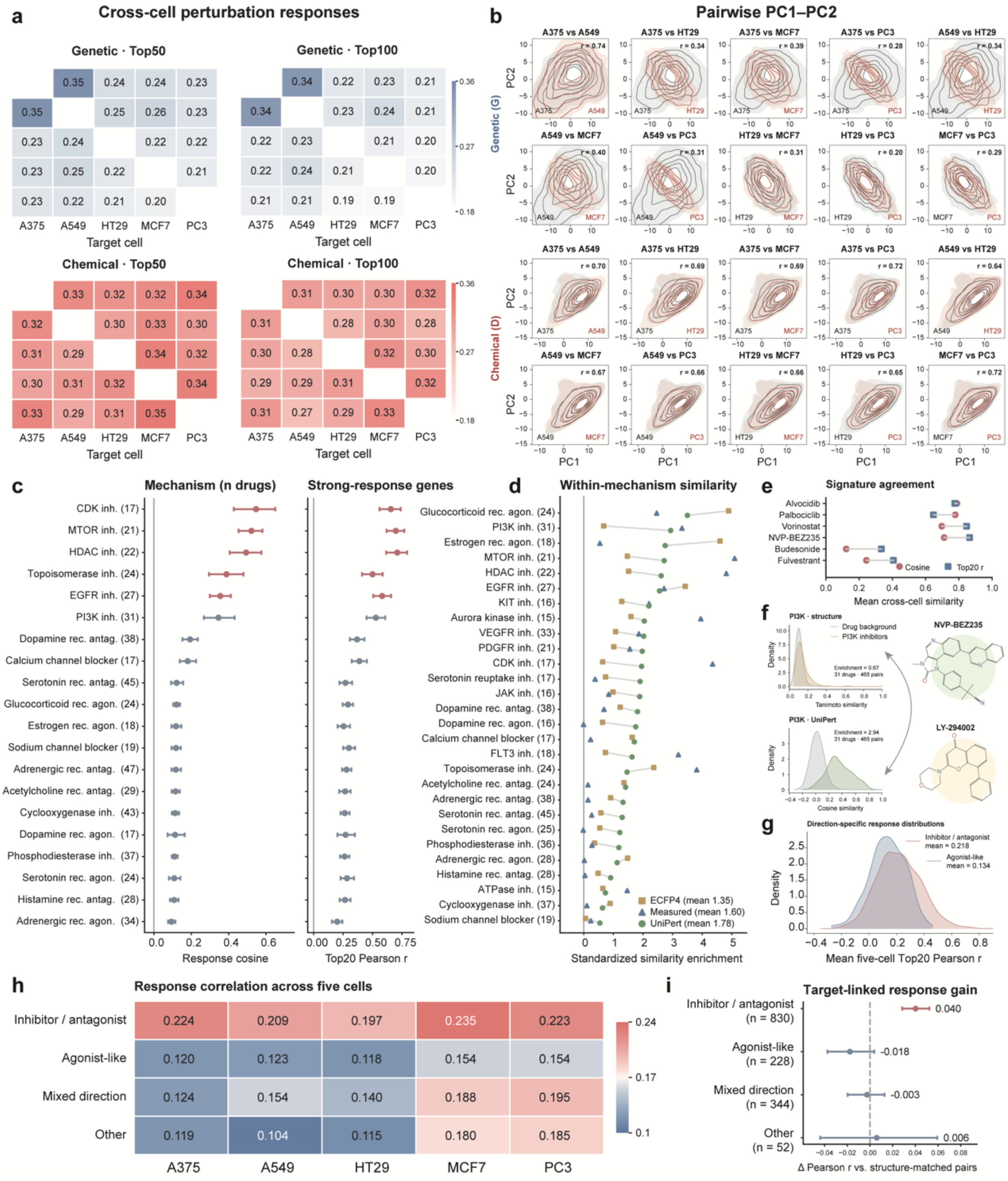


**Extended Data Fig. 1 | Cross-context response conservation and genetic–chemical relationships.** **a**, Mean cross-cell Pearson correlations on the 50 or 100 genes with largest absolute responses in the source cell (rows); columns denote targets. Genetic and chemical analyses include 4,156 and 3,220 shared perturbations, respectively. **b**, Pairwise PC1–PC2 density contours in a common basis fitted without MCF7. Annotated *r* is the Pearson correlation of matched-perturbation PC1 scores. **c**, Cross-cell cosine similarity and source-defined Top20 Pearson correlations by primary mechanism of action (MoA); points and whiskers show drug means and 95% bootstrap intervals. Parentheses indicate drug numbers. **d**, Within-MoA similarity enrichment relative to the drug-library background, in background standard-deviation units, for ECFP4, measured responses and UniPert. Classes can overlap; each contains at least 15 drugs. **e**, Mean cross-cell cosine similarity and Top20 correlation for six example drugs. **f**, ECFP4 Tanimoto and UniPert cosine similarities among 31 PI3K inhibitors (465 pairs) compared with the drug-library background, with example structures. **g**,**h**, Correlations between measured genetic responses and responses to drugs targeting the perturbed genes, stratified by intervention direction; Top20 genes are defined by genetic responses. Distributions in **g** average each pair across five cells; **h** shows cell means. **i**, Drug-level correlation gain over structure-matched non-target pairs. Points and whiskers show means and 95% intervals from 2,000 drug bootstraps; *n* denotes drugs. All five-cell analyses use A375, A549, HT29, MCF7 and PC3.

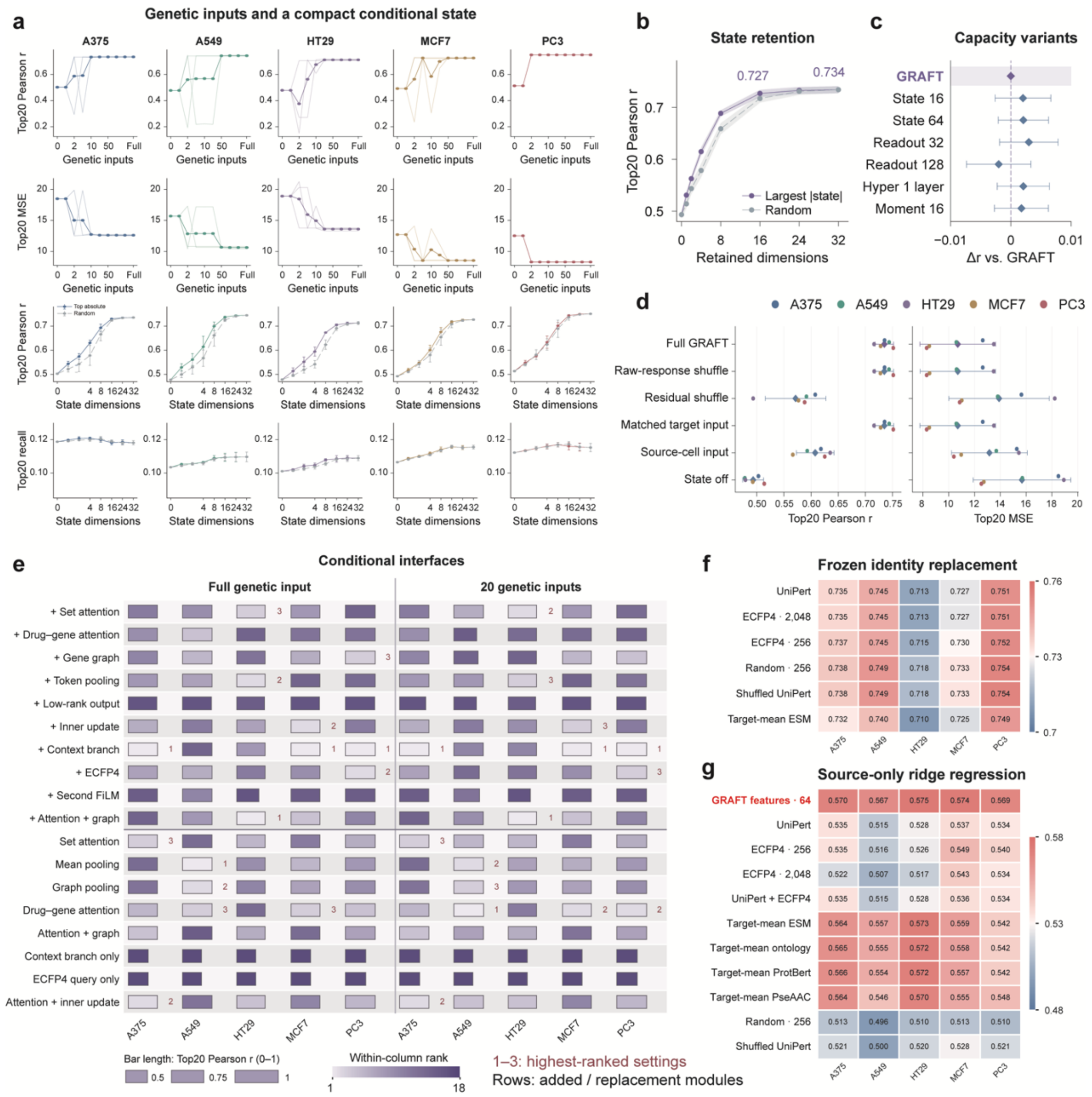


**Extended Data Fig. 2 | Genetic conditioning, capacity controls and representation probes. a**, Prediction accuracy versus target genetic-input number (top two rows) or retained state dimensions (bottom two rows) for five held-out cell lines. Thin genetic-budget curves denote three training seeds; thick curves are means. State-retention points and whiskers show mean ± s.d. across seeds, retaining largest-magnitude or random coordinates of the frozen 32-dimensional state. **b**, State retention averaged across five cells after seed averaging; shading denotes s.e.m. **c**, Accuracy differences for retrained capacity variants relative to their matched GRAFT reference; diamonds and whiskers show five-cell means and 95% *t*-intervals after averaging three seeds per cell. **d**, Frozen-model genetic-input interventions. Raw-response shuffling recomputes residuals against unchanged source references; residual shuffling disrupts residual–identity correspondence. Matched target and source-cell inputs use the same genetic-perturbation intersection. Colored points denote cell means; diamonds and whiskers show five-cell means and 95% *t*-intervals. **e**, Added modules (above divider) and replacement interfaces (below), evaluated with full or 20 genetic inputs; bar length denotes seed-mean correlation, color denotes within-column rank and numbers identify ranks 1–3. **f**, Frozen drug-identity replacement after linear projection into the native identity space, with source responses and genetic inputs held fixed. **g**, Residual ridge models fitted on non-query target drug measurements to correct source-mean predictions. Heatmaps show three-seed means; protein-derived probes use 364 queries, versus 644 for other representations. Of the full 644-query benchmark, 641 have defined correlations.

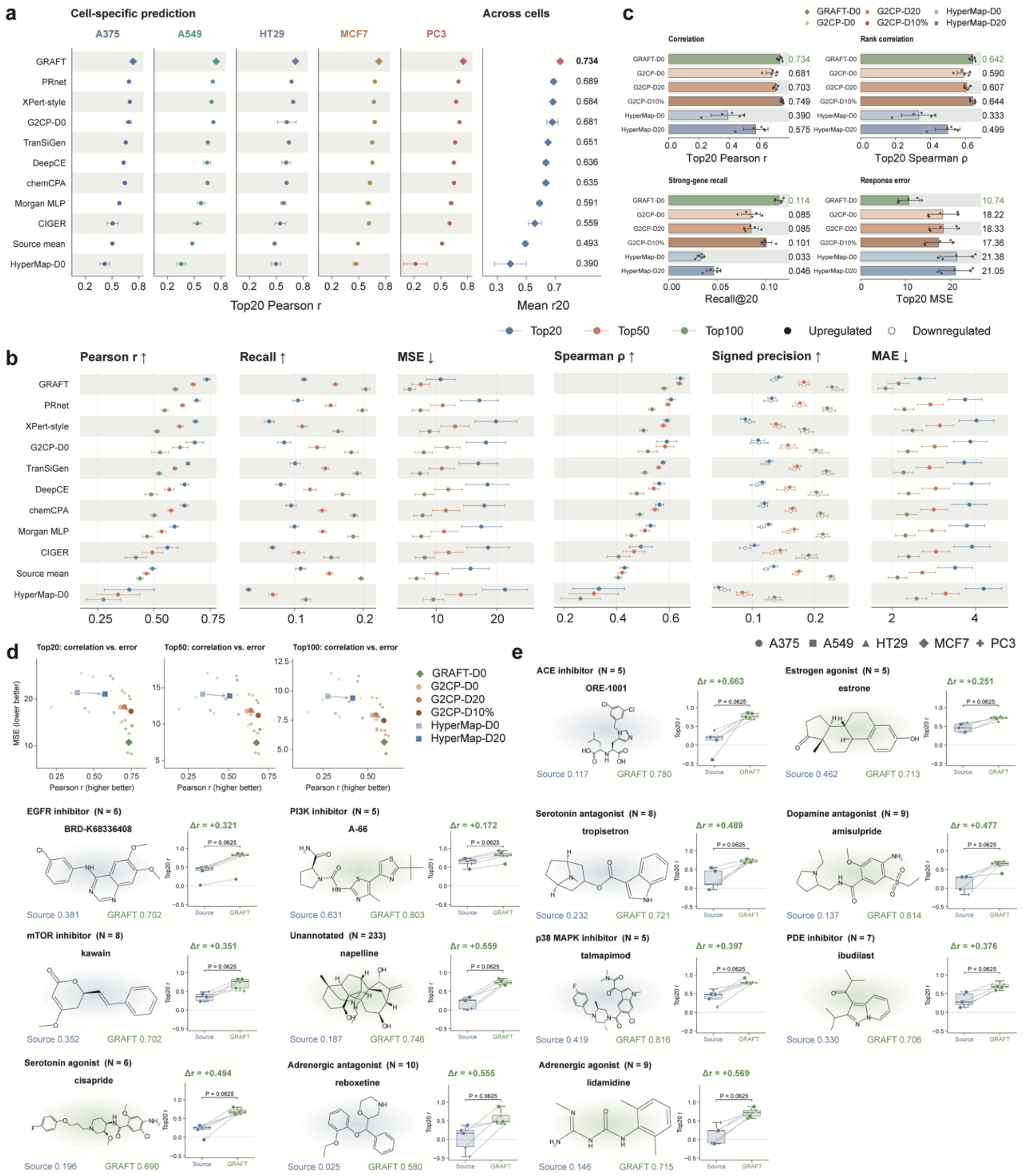


**Extended Data Fig. 3 | Detailed LINCS prediction benchmarks and pharmacological examples. a**, Top20 Pearson correlations across five held-out cell lines and 11 D0 methods. Cell-specific points show three-seed means ± s.d.; the right panel shows five-cell means ± s.d. **b**, Six prediction metrics across Top20, Top50 and Top100 response windows (Methods). Points and whiskers show five-cell means ± s.d. after seed averaging; filled and open symbols denote upregulated- and downregulated-gene precision. **c**, Correlation, rank correlation, strong-response gene recall and error for GRAFT-D0, G2CP and HyperMap under different target-drug budgets. Bars and whiskers show five-cell means ± s.d.; black points denote cell means. D0 uses no target drug measurements; D20 uses 20 non-query target drugs; D10% uses 10% of the target drug catalog (339–768 drugs), drawn from the non-query pool. **d**, Correlation–error trade-offs across response windows; small points denote cells, large symbols denote macro means, and arrows connect increasing within-model budgets. **e**, Example compounds with the largest five-cell mean GRAFT-minus-source correlation gain within each displayed primary-MoA group. *N* denotes candidate drugs, whereas the paired sample size is five cell lines. Shapes identify cells and lines join matched observations. Boxes show medians and interquartile ranges; whiskers extend to 1.5 times the interquartile range. Displayed *P* values are from unadjusted two-sided exact paired sign-flip tests (32 assignments); none remained significant after Holm correction across 641 eligible compounds. Benchmark evaluations use 644 fixed query drugs, with 641 defined correlation profiles.

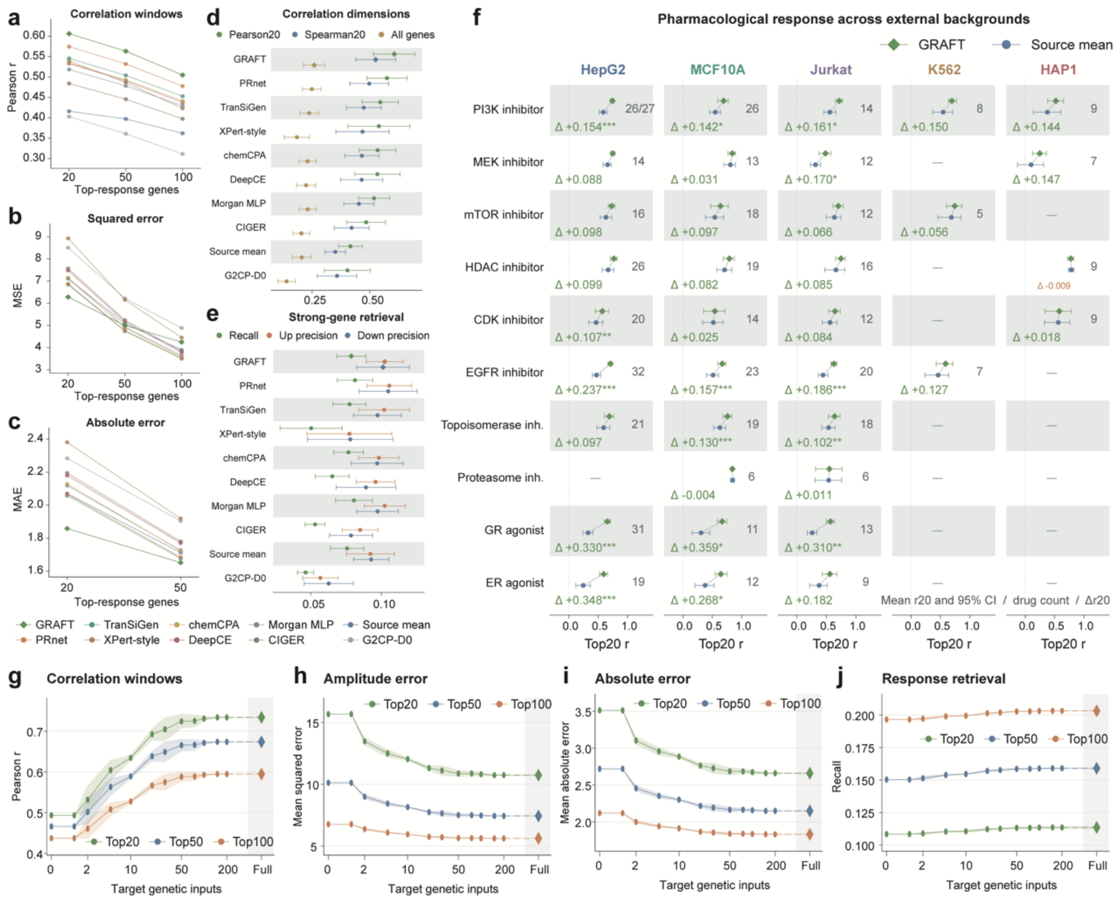


**Extended Data Fig. 4 | Extended external benchmarks and genetic-input requirements.** **a–c,** Pearson correlations (a), mean squared error (MSE; b) and mean absolute error (MAE; c) across strong-response gene windows for ten D0 methods. Windows select genes by absolute measured response. Curves show equally weighted means across five external contexts after averaging three training seeds within each context. **d,e,** Top20 Pearson and Spearman correlations and all-gene Pearson correlation (d), and Top20 recall and upregulated/downregulated gene precision (e). Points and error bars indicate five-context means ± s.d. **f,** Class-stratified Top20 Pearson correlations for GRAFT and the source mean. Points and intervals show drug means and 95% drug-bootstrap confidence intervals after averaging seeds. Counts indicate valid paired drugs, with total class counts added when different; dashes indicate fewer than five drugs. Δ denotes GRAFT minus source mean. Asterisks indicate two-sided paired Wilcoxon tests across drugs, with Holm correction across the 36 displayed context–class comparisons: $P < 0.05$ (*), $P < 0.01$ (**) and $P < 0.001$ (***). MoA annotations may overlap. **g–j,** Internal five-cell prediction performance as target genetic inputs increase: Pearson correlation (g), MSE (h), MAE (i) and recall (j), using Top20, Top50 or Top100 genes. Curves and shading show means ± s.d. across three seeds after averaging three input draws per seed and then the five target contexts. Diamonds indicate full-input results.

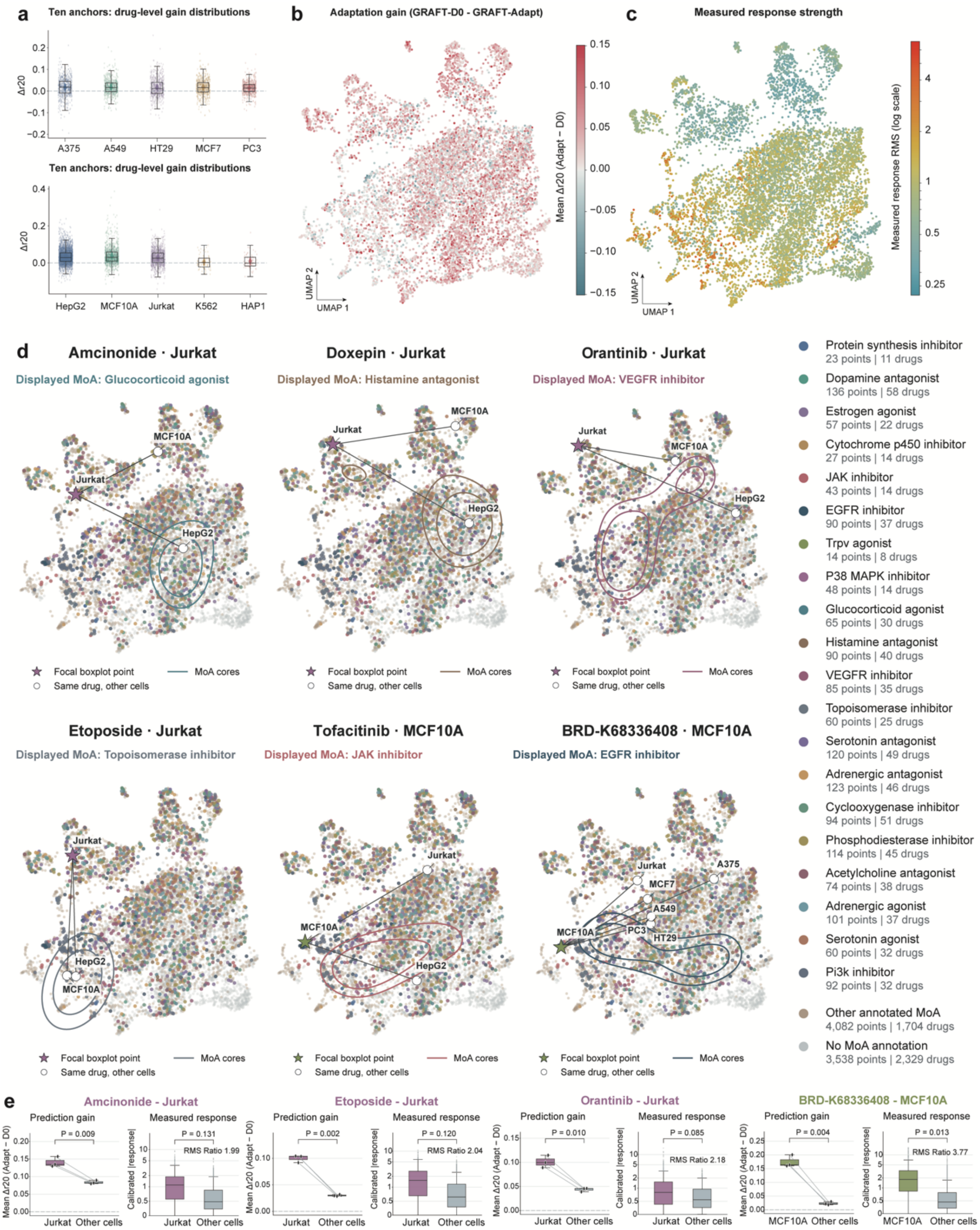
a
Ten anchors: drug-level gain distributions
Δr20
A375
A549
HT29
MCF7
PC3
Ten anchors: drug-level gain distributions
Δr20
HepG2
MCF10A
Jurkat
K562
HAP1
b
Adaptation gain (GRAFT-D0 - GRAFT-Adapt)
Mean Δr20 (Adapt − D0)
UMAP 1
UMAP 2
c
Measured response strength
Measured response RMS (log scale)
UMAP 1
UMAP 2
d
Amcinonide · Jurkat
Displayed MoA: Glucocorticoid agonist
Doxepin · Jurkat
Displayed MoA: Histamine antagonist
Orantinib · Jurkat
Displayed MoA: VEGFR inhibitor
Focal boxplot point
Same drug, other cells
MoA cores
Etoposide · Jurkat
Displayed MoA: Topoisomerase inhibitor
Tofacitinib · MCF10A
Displayed MoA: JAK inhibitor
BRD-K68336408 · MCF10A
Displayed MoA: EGFR inhibitor
Protein synthesis inhibitor
23 points | 11 drugs
Dopamine antagonist
136 points | 58 drugs
Estrogen agonist
57 points | 22 drugs
Cytochrome p450 inhibitor
27 points | 14 drugs
JAK inhibitor
43 points | 14 drugs
EGFR inhibitor
90 points | 37 drugs
Trpv agonist
14 points | 8 drugs
P38 MAPK inhibitor
48 points | 14 drugs
Glucocorticoid agonist
65 points | 30 drugs
Histamine antagonist
90 points | 40 drugs
VEGFR inhibitor
85 points | 35 drugs
Topoisomerase inhibitor
60 points | 25 drugs
Serotonin antagonist
120 points | 49 drugs
Adrenergic antagonist
123 points | 46 drugs
Cyclooxygenase inhibitor
94 points | 51 drugs
Phosphodiesterase inhibitor
114 points | 45 drugs
Acetylcholine antagonist
74 points | 38 drugs
Adrenergic agonist
101 points | 37 drugs
Serotonin agonist
60 points | 32 drugs
Pi3k inhibitor
92 points | 32 drugs
Other annotated MoA
4,082 points | 1,704 drugs
No MoA annotation
3,538 points | 2,329 drugs
e
Amcinonide - Jurkat
Prediction gain
Measured response
P = 0.009
P = 0.131
RMS Ratio 1.99
Etoposide - Jurkat
P = 0.002
P = 0.120
RMS Ratio 2.04
Orantinib - Jurkat
P = 0.010
P = 0.085
RMS Ratio 2.18
BRD-K68336408 - MCF10A
P = 0.004
P = 0.013
RMS Ratio 3.77
Mean Δr20 (Adapt − D0)
Calibrated |response|
Jurkat
MCF10A
Other cells

**Extended Data Fig. 5 | Adaptation gains and context-dependent measured drug responses. a,** Drug-level Top20 correlation gains with ten target-drug anchors across five internal and five external contexts. Each point is a drug mean across training seeds and anchor draws; diamonds indicate means across drugs. **b,c,** Fixed UMAP of 9,136 measured drug–context profiles from eight contexts, colored by mean adaptation gain (GRAFT-Adapt minus GRAFT-D0; b) or measured-response RMS on a logarithmic scale (c). Coordinates were derived from centered, normalized measured responses; coloring did not determine the embedding. **d,** Six illustrative drug–context comparisons in the same measured-response space. Stars mark focal contexts; open circles show the same drug in other measured contexts; connecting lines identify the same drug. Contours indicate MoA reference densities after excluding case drugs. Display colors assign one annotation per point; density estimates retain all matching annotations. **e,** Focal-versus-other-context comparisons for four examples. Gain plots show three paired training-seed means, each averaged across 30 anchor draws; other contexts are equally weighted within each seed. Displayed *P* values are unadjusted, two-sided paired *t* tests ($n$ = 3 seed pairs). Response plots show absolute measured responses for 976 genes per context, calibrated by median RMS across 56 shared reference drugs. Response *P* values are two-sided empirical probabilities for whole-profile RMS specificity relative to other drugs measured in the same contexts; genes are not independent replicates. Boxes in **a,e** indicate medians and interquartile ranges; whiskers extend to observations within 1.5 times the interquartile range.

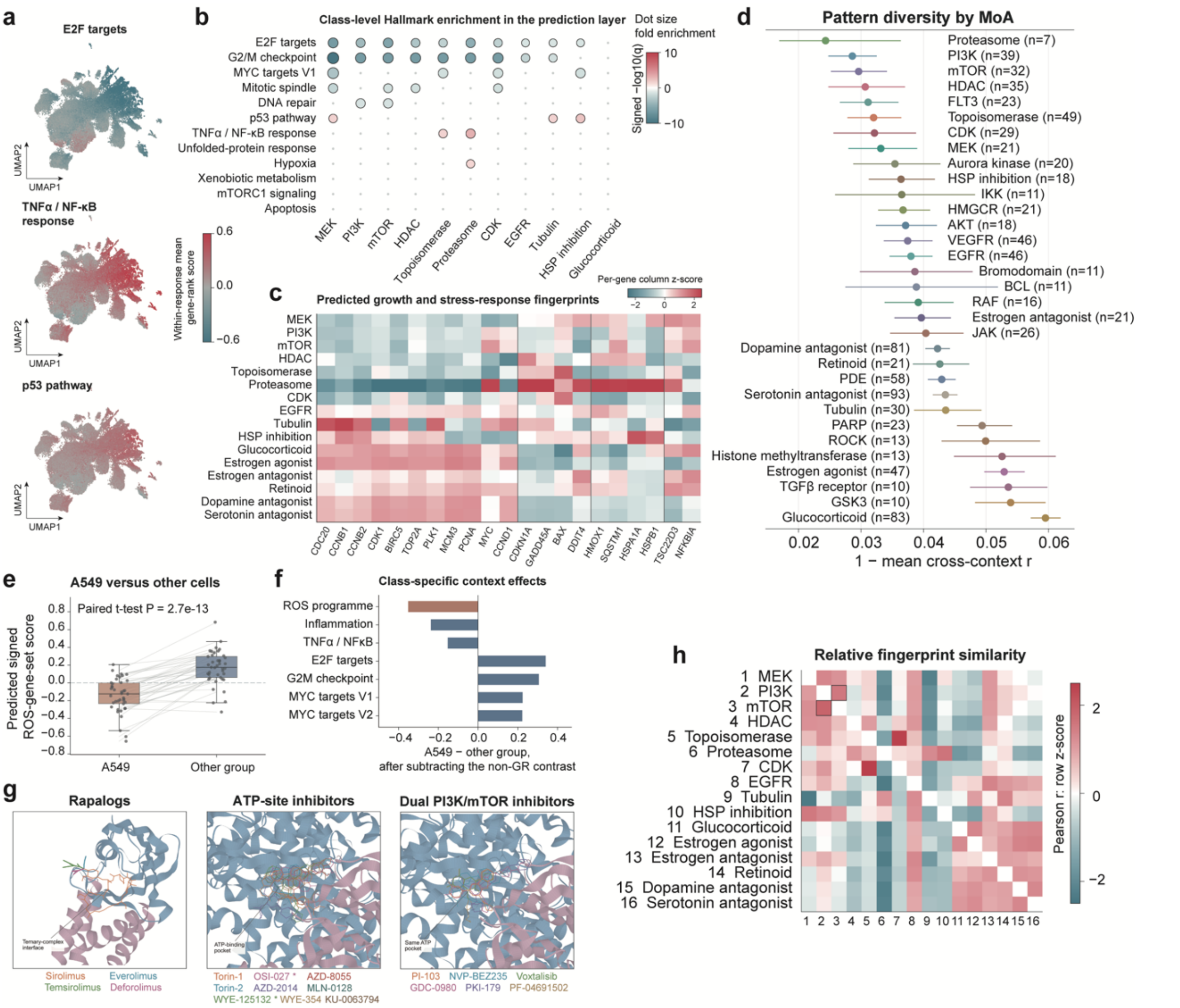


**Extended Data Fig. 6 | Predicted functional programs, context effects and mTOR structural references. a,** Prediction-space UMAP colored by within-profile mean gene-rank scores for E2F, TNFα/NF-κB and p53 programs. **b,** Hallmark overrepresentation among the 50 highest or lowest genes in context-averaged class predictions, using the 976-gene background. Bubble area indicates fold enrichment; color indicates direction-signed −log10(*q*). Hypergeometric *P* values were Benjamini–Hochberg corrected across 21,952 tests; colored bubbles have $q < 0.05$. **c,** Predicted growth- and stress-related gene fingerprints, averaging class means across 13 contexts and standardizing each gene across classes. **d,** Class-specific pattern diversity, defined as one minus the mean cross-context Pearson correlation per chemical. Points and intervals show class means and 95% confidence intervals from 5,000 chemical-entity bootstrap resamples; *n* denotes annotated entities. **e,** Paired predicted ROS-gene-set scores for 41 glucocorticoid connectivity groups in A549 versus their means across A375, HT29, MCF7 and PC3; two-sided paired *t* test. Boxes indicate median and interquartile range; whiskers extend to 1.5 times the interquartile range. **f,** Mean signed gene-set-score differences between A549 and the other four contexts for glucocorticoids, after subtracting the corresponding non-glucocorticoid differences. **g,** Structural references for four rapalogs at the FKBP12–FRB interface (PDB 1FAP), and computed poses of nine ATP-site and six dual PI3K/mTOR inhibitors in the mTOR ATP pocket (PDB 4JSX). Rapalog extensions are illustrative core-aligned overlays, not docking predictions. **h,** Relative class-fingerprint similarities across 976 genes, with shared class members excluded pairwise; colors show off-diagonal row-standardized Pearson correlations. Boxes highlight PI3K–mTOR similarity. All response analyses use predictions: GRAFT-Adapt in ten contexts and D0 in RPE1, neurons and LUHMES.
